# The genetic driver of Acute Necrotizing Encephalopathy, *RANBP2*, regulates the inflammatory response to Influenza A virus infection

**DOI:** 10.1101/2025.03.23.644734

**Authors:** Sophie Desgraupes, Adrien Decorsière, Suzon Perrin, Benoît Gouy, Yifan E Wang, Alexander F. Palazzo, Sandie Munier, Nathalie J. Arhel

## Abstract

Influenza virus infections can cause severe complications such as Acute Necrotizing Encephalopathy (ANE), which is characterised by rapid onset pathological inflammation following febrile infection. Heterozygous dominant mutations in the nucleoporin RANBP2/Nup358 predispose to influenza-triggered ANE1. The aim of our study was to determine whether RANBP2 plays a role in IAV-triggered inflammatory responses. We found that the depletion of RANBP2 in a human airway epithelial cell line increased IAV genomic replication by favouring the import of the viral polymerase subunits, PB1, PB2 and PA following viral transcription and translation. Additionally, RANBP2 knockdown enhanced the cytoplasmic export of viral RNA (vRNA) and disrupted segment stoichiometry, which associated with elevated production of the pro-inflammatory chemokines CXCL8, CXCL10, CCL2, CCL3 and CCL4 in human primary macrophages. Using CRISPR-Cas9 knock-in for the ANE1 disease variant RANBP2-T585M, we further demonstrate that this point mutation causes a loss-of-localisation phenotype that excludes RANBP2 from the nuclear envelope, which phenocopies RANBP2 knockdown by increasing IAV replication and driving pro-inflammatory cytokine expression following infection. Together, our results reveal that RANBP2 regulates influenza RNA replication and nuclear export, thereby restraining virus-induced hyperinflammation, and further suggest that ANE1 pathogenesis results from the impaired localisation of RANBP2 at the nuclear envelope.

## INTRODUCTION

Influenza viral infections are responsible for seasonal epidemics with respiratory manifestations but can also lead to severe complications. Among these, Acute Necrotizing Encephalopathy (ANE), also called ANE of Childhood (ANEC), is a severe reaction to febrile infection that is characterized by rapid progression and very poor prognosis (Lee et al., 2023; Sakuma et al., 2025; Wu et al., 2015; Mizuguchi et al., 2007). Influenza viruses are thought to be responsible for ∼50% of all ANE episodes and are associated with high morbidity and mortality (Bartolini et al., 2024; Jiang et al., 2022; Mizuguchi et al., 2007; Neilson et al., 2003; Chatur et al., 2022; Bashiri et al., 2020). Although the pathogenesis is unclear, a hyperinflammatory response triggered by the virus is proposed as the underlying mechanism (Levine et al., 2020; Mizuguchi et al., 2007; Sugaya, 2002), and early anti-inflammatory therapy may help mitigate the disease (Chatur et al., 2022; Koh et al., 2019; Okumura et al., 2009).

Influenza viruses belong to the *Orthomyxoviridae* family and are classified into types, Influenza A and B being the most prevalent ones associated with seasonal flu cases. Influenza A is further divided into subtypes based on their surface glycoproteins, haemagglutinin (HA) and neuraminidase (NA). The H1N1 and H3N2 strains are responsible for major outbreaks and epidemics, and in the 2024-2025 winter season, there has been an unprecedented surge in H1N1 cases, leading to an unusual spike in hospitalizations and ANE in children (Bartolini et al., 2024).

The Influenza A virus (IAV) is an enveloped particle containing 8 negative-sense single-stranded viral RNA (vRNA) gene segments, each packaged by nucleoprotein (NP) into a viral ribonucleoprotein (vRNP) complex, and associated with the viral polymerase complex (PB1, PB2, and PA). Unlike most RNA viruses, IAV replicates in the nucleus and hijacks the nuclear pore complex (NPC) machinery for four critical steps of its replication cycle. (i) After endocytosis and uncoating, vRNPs are trafficked to the nucleus where the viral polymerase transcribes the viral genome. (ii) Capped and polyadenylated mRNAs are exported into the cytoplasm for translation by cytoplasmic ribosomes. (iii) The newly translated viral proteins, including NP, PB1, PB2 and PA, are imported back into the nucleus, which allows the viral polymerase to replicate the viral genome into complementary RNAs (cRNA) and genomic vRNA. (iv) Lastly, new vRNPs, assembled with M1 and NS2/NEP, are exported back from the nucleus for incorporation into budding virions (Dou et al., 2018; Carter and Iqbal, 2024; Fodor and Te Velthuis, 2020; Zhu et al., 2023).

Many viral and host factors are thought to contribute to severe influenza, and genetic susceptibility particularly in genes that regulate innate signalling and antiviral response, has been reported (Mettelman and Thomas, 2021; Clohisey and Baillie, 2019; Gounder and Boon, 2019). However, the mechanisms that drive severe influenza-triggered ANE in previously healthy children are not known. One clue to understanding the pathogenesis of influenza-associated ANE comes from genetic susceptibility in familial, recurrent cases, known as ANE1 (or ADANE) (Neilson et al., 2003). Patients with ANE1 present with autosomal dominant missense mutations in *RANBP2* and have a significantly increased lifetime risk of developing ANE (Neilson et al., 2009). Mutations in *RANBP2* increase the risk of relapse and recurrence (Levine et al., 2020; Chatur et al., 2022) and may associate with greater morbidity and mortality.

RANBP2 (Nup358) is a cytoplasmic fibril (CF) nucleoporin that regulates nucleocytoplasmic transport (NCT) by interacting with nuclear transport receptors such as Karyopherin α/β and CRM1/Exportin 1 (Yokoyama et al., 1995; Hutten et al., 2008). Additional functions have been attributed to RANBP2 at the NPC, many of which are linked to its C-terminal domain (CTD) E3 SUMO ligase activity, and away from the NPC, namely at annulate lamellae, mitochondria-endoplasmic reticulum junctions and in the nucleus (Desgraupes et al., 2023). Moreover, at least 4 protein isoforms have been identified for RANBP2, however, it is not known if these have distinct functions or subcellular localisations (Desgraupes & Arhel, 2025). The predominant ANE1-associated mutation (c.C1754T on the coding DNA reference sequence, leading to p.T585M at the protein level), and most other pathogenic mutations in RANBP2, all cluster in the N-terminal domain (NTD), which is responsible for anchoring to NPCs (Jiang et al., 2022; Joseph and Dasso, 2008). This suggests that the localization of RANBP2 to CF of NPC may be physiologically essential, although no changes in localisation could be demonstrated following the overexpression of disease variants (Shen et al., 2021; Bley et al., 2022).

Previous work showed that RANBP2 can modulate viral infection of several viruses, such as Adenovirus (Carlon-Andres et al., 2020), Herpes simplex virus type 1 (HSV-1) (Copeland et al., 2009; Hofemeister and O’Hare, 2008) and Human immunodeficiency virus type 1 (HIV-1) (Bichel et al., 2013; Di Nunzio et al., 2012; Schaller et al., 2011; Zhang et al., 2010), however no work has uncovered a role for RANBP2 in Influenza virus infection. In fact, although some studies showed that NPC components can impact the replication and transcription of Influenza virus, such as Nup62, Nup98 and Nup153, RANBP2 was not identified as a host factor for Influenza virus (Khanna et al., 2024; Watanabe et al., 2010; Munier et al., 2013).

Previous work also showed that RANBP2 can modulate innate immune signalling, however this activity was consistently attributed to its E3 SUMO ligase activity or Cyclophilin-homology motif in the CTD (Maarifi et al., 2018; Portilho et al., 2016; Shen et al., 2021; Li et al., 2023), and no work has yet demonstrated a role for the NTD where ANE1 mutations are clustered.

The aim of our study was to determine how RANBP2 contributes to or regulates inflammatory responses to infection by Influenza virus. We show that RANBP2 impacts the replication of genomic vRNAs of IAV into cRNA by controlling the re-import of newly translated viral polymerase complex. In addition, RANBP2 knockdown enhances cytoplasmic vRNA export and disrupts segment stoichiometry, thereby enhancing IAV-triggered inflammation. Notably, the predominant ANE1-associated variant T585M causes relocation of RANBP2 away from the nuclear envelope, which phenocopies knockdown of RANBP2 by increasing vRNA replication and export, and triggering hyperinflammation, which could provide a first clue in understanding the pathogenesis of ANE.

## RESULTS

### Knockdown of RANBP2 stimulates non-productive Influenza vRNA replication in infected cells

To investigate the role of RANBP2 in IAV infection, RANBP2-knockdown (RANBP2-KD) was induced in A549 human alveolar cells by lentiviral vector (LV) transduction before infection with A/WSN/1933 (H1N1) virus at MOI 0.5. Supernatants were harvested at 10 h post-infection (hpi) to investigate the production of infectious viral particles by TCID50 titration. No differences were observed between the control and RANBP2-KD cells, suggesting that RANBP2 had no effect on the production of infectious particles **(Figure 1A, Figure S1A-B)**.

**Figure 1:**
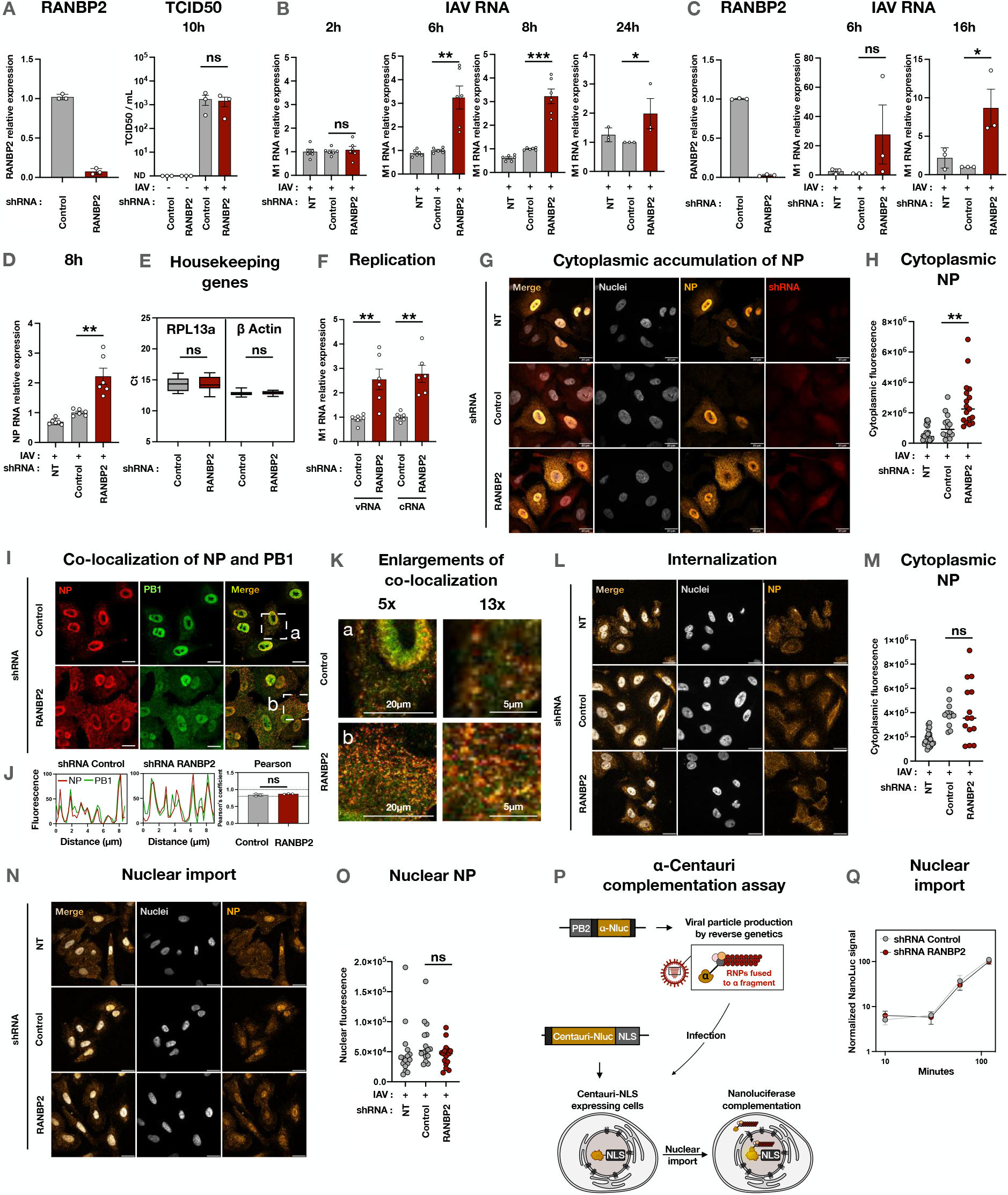
RANBP2 knockdown stimulates non-productive Influenza vRNA replication without affecting the initial vRNP nuclear import. **(A)** A549 cells transduced with LV shRNA-control or shRNA-RANBP2 were infected with IAV (A/WSN/1933; MOI 0.5). RANBP2 transcript levels were quantified by RT-qPCR and normalized on RPL13a housekeeping gene and on the control condition. Viral titers in supernatants at 10 hpi were determined by TCID50 on MDCK cells. **(B)** A549 cells or **(C)** THP-1 cells were transduced with LV shRNA-control or shRNA-RANBP2 or non-treated (NT) then infected with influenza A virus (A/WSN/1933; MOI 0.5). At the indicated time points, cells were lysed and total intracellular RNAs were extracted. Viral RNAs encoding the Matrix 1 (M1) protein were amplified by RTqPCR. **(D)** Nucleoprotein-encoding (NP) viral RNAs were amplified by RTqPCR in the A549 infected cells. **(E)** Housekeeping gene expression was not affected by the knockdown. n=18 and n=12 independent experiments for RPL13a and β-Actin, respectively. **(F)** Transduced A549 cells were infected with IAV (MOI 0.5) and total intracellular RNAs were extracted at 6 hpi. Strand-specific reverse transcription (RT) was performed with tagged-primers to amplify separately the genomic viral RNA (vRNA) and the complementary viral RNA (cRNA). M1-encoding viral RNAs were amplified by RTqPCR. Results are normalized on the control condition. **(G)** NP staining in transduced and infected A549 cells at 8 hpi. The images from 4 additional representative fields are provided for each condition in Figure S3. **(H)** Quantification of cytoplasmic NP fluorescence. Each dot represents the cytoplasmic fluorescence of a single cell, normalized by its area. **(I)** Co-staining of NP and PB1 in transduced and infected A549 cells at 8 hpi. **(J)** Colocalization curves and Pearson’s correlation coefficients between PB1 and NP signals. **(K)** Enlargements of colocalization fields from panel I. **(L)** NP staining in transduced and infected A549 cells at 1 hpi (MOI 4). **(M)** Quantification of cytoplasmic NP fluorescence. Each dot represents the cytoplasmic fluorescence of a single cell, normalized by its area. **(N)** NP staining in transduced and infected A549 cells at 2 hpi (MOI 4). **(O)** Quantification of nuclear NP fluorescence. Each dot represents the nuclear fluorescence of a single cell. (P) Experimental strategy of the Alpha-Centauri assay. IAV (A/WSN/1933) particles containing the PB2 protein fused to a fragment of the NanoLuc (IAV-α) were produced by reverse genetics. A plasmid containing a nuclear localization signal (NLS) fused to the other fragment of the NanoLuc (Cen-NLS) was generated and transfected into A549 cells. (Q) Cen-NLS expressing cells were infected with IAV-α (MOI 4) and the luminescence was measured to assess the NanoLuc complementation upon nuclear import of the viral ribonucleoproteins. NanoLuc complementation results are normalized on the last time point (n = 3 independent experiments). In qPCR quantifications (panels A, B, C, D, F), dots represent independent biological replicates, each corresponding to the mean of technical triplicates normalized to a housekeeping gene (ΔCt). Data are plotted relative to the control condition (2^-ΔΔCt) and shown as mean +/− SEM. Confocal images (panels G, I, K, L, N) are representative of n = 3 independent experiments. All scale bars: 20µm. Imaging quantifications (panels H, J, M, O) were performed using Fiji and show individual cells (n = 3 independent experiments). Statistical analyses were performed on biological replicates only. Two-tailed unpaired (panels A, B, E, F, H, J, M, O) or paired (C-D) Student’s t tests were performed using Prism 10. *P < 0.05, **P < 0.01, ***P < 0.001, ns: non-significant.

However, an unexpected phenotype was observed for viral RNA quantification in both A549 and the monocytic cell line THP-1. Infection kinetics were performed and viral RNA levels were determined at early (2 h) and late (6, 8, 16 and 24 h) time points using qPCR primers recognizing M1 on segment 7. In A549 cells **(Figure 1B)** and in the monocytic cell line THP-1 **(Figure 1C, Figure S1C)**, a significant increase of M1 RNA was induced in RANBP2-KD cells. The increase in viral RNA was confirmed in A549 cells with another IAV segment, encoding the NP **(Figure 1D)**, and appeared to be specific since no increase was observed for any of the housekeeping genes that were measured **(Figure 1E)**. Together, these data suggested that RANBP2 is involved in controlling Influenza viral RNA synthesis but that this does not translate into increased productive infection.

Because different types of viral RNAs are produced during the IAV life cycle, we investigated which RNAs are impacted by the depletion of RANBP2. Sequence-tagged RT primers were used to specifically recognize the negative-sense genomic vRNA sequence or the positive-sense cRNA sequence in order to specifically reverse transcribe (RT) each type of viral RNA **(Figure S2A)**. First, the specificity of each primer was validated by amplifying only vRNAs after vRNA-specific RT, or only cRNAs after cRNA-specific RT, while not amplifying any target in non-infected cells **(Figure S2B)**. In RANBP2-KD cells, we observed a significant increase of both vRNAs and cRNAs **(Figure 1F)**, thus showing that RANBP2 is involved in IAV RNA replication. Although this was unexpected given the absence of phenotype in terms of viral production, the marked increase in vRNA and cRNA levels suggested an enhancement in viral genome replication.

Having observed an increase in genomic vRNAs, we tested if this correlated with more newly produced vRNPs. As previously, A549 cells were transduced with control- or RANBP2-shRNAs then infected with IAV (MOI 0.5). At 8 hpi, the distribution of NP was investigated by immunofluorescence. Although export is ongoing in every condition, RANBP2-KD cells showed a striking increase in cytoplasmic NP compared to control **(Figure 1G-H, Figure S3)**. This phenotype was confirmed using a previously published RANBP2-targeting shRNA (Di Nunzio et al., 2012) and only in cells expressing the shRNA (**Figure S4A-D**). In order to determine if signal corresponded to vRNPs or free NP in the cytoplasm, we performed co-stainings with other components of vRNPs (i.e. PB1 and PA). We observed a co-localisation of cytoplasmic NP with PB1 and with PA, with a Pearson coefficient >0.8 **(Figure 1I-K, Figure S5A-D)**, indicating that RANBP2-KD induces an increase in vRNPs exported to the cytoplasm.

### Increased IAV replication does not result from enhanced viral entry into cells or into the nucleus

To determine how vRNA is increased in RANBP2-depleted cells, we first tested whether RANBP2 regulates IAV infection by modulating viral entry into cells and/or nuclei, thereby increasing the quantity of IAV RNA to be copied.

To assess whether RANBP2 regulates cell or nuclear entry, control or RANBP2-KD cells were infected with IAV (A/WSN/33) at high MOI of 4, in order to increase signal detection during early steps of the viral cycle, and NP localization was studied by immunofluorescence at 1 hpi to assess internalization, and at 2 h to monitor NP nuclear entry. Results indicate that depletion of RANBP2 from A549 cells affects neither internalization **(Figure 1L-M)** nor nuclear import **(Figure 1N-O)**.

To further validate these findings, we adapted an Alpha-Centauri protein-complementation assay previously developed to monitor the nuclear import of HIV-1 (Fernandez et al., 2021). Briefly, the NanoLuc protein was expressed as two unequal non-luminescent fragments. The smaller fragment was fused to PB2 in IAV particles (IAV-α), while the larger fragment (Centauri) was fused to a NLS and expressed in A549 cells by LV transduction (Cen-NLS) **(Figure 1P, Figure S6A-B)**. Upon vRNP nuclear import, the two fragments reconstitute a functional NanoLuc, generating a measurable signal upon addition of substrate **(Figure S6C)**. No significant differences in luminescence were detected between control and RANBP2-KD cells at 1 and 2 hpi, confirming that RANBP2 is not essential for vRNP nuclear import **(Figure 1Q, Figure S6D)**, thus demonstrating that RANBP2 does not regulate IAV vRNP nuclear import.

### RANBP2 knockdown facilitates the nuclear import of the neo-synthesised viral polymerase complex proteins PB1, PB2 and PA

Since RANBP2 does not impact the quantity of template available for replication, we investigated whether it modulates the quantity or localisation of the polymerase complex subunits. First, the total levels of PB1, PB2, PA and NP were assessed by western blot at 8 hpi, however, no differences were observed between RANBP2-KD cells and control cells **(Figure 2A-B)**, suggesting that RANBP2 does not impact the translation of the viral polymerase complex.

**Figure 2:**
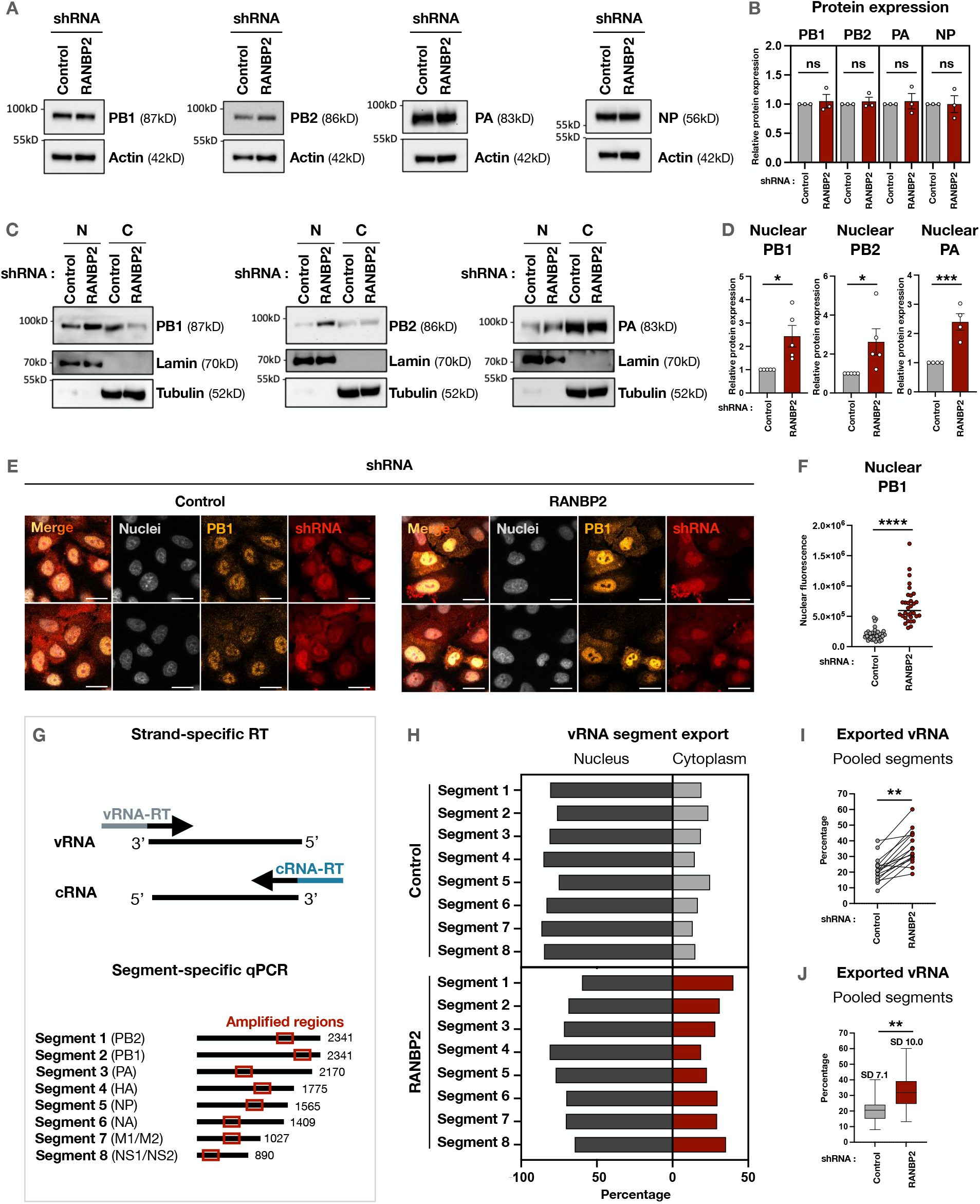
RANBP2 knockdown enhances nuclear reimport of the viral polymerase complex proteins PB1/PB2/PA and cytoplasmic vRNP accumulation with disrupted segment stoichiometry. **(A)** A549 cells transduced with LV shRNA-control or shRNA-RANBP2 were infected with IAV (A/WSN/1933; MOI 4). At 6 hpi, PB1, PB2, PA and NP protein levels were determined by western blot. **(B)** Band intensities from n =3 independent experiments were quantified using ImageLab. Values are normalized on the actin signal and to the control condition. **(C)** Transduced A549 cells were treated with KPT-330 for 2 h, infected with IAV (MOI 0.5) and subjected to nucleo-cytoplasmic fractionation at 6 hpi. PB1, PB2 and PA levels were determined by western blot (N: nucleus, C: cytoplasm). Lamin and tubulin protein levels were investigated to verify the efficacy of the fractionation. **(D)** Nuclear band intensities from n =5 independent experiments were quantified using ImageLab. Results are normalized on the lamin signal and to the control condition. **(E)** Transduced A549 cells were infected with IAV (MOI 0.5) in the absence of KPT-330 and stained for PB1 at 6 hpi. Scale bar: 20µm. **(F)** Nuclear PB1 fluorescence was quantified using Fiji. Each dot represents the nuclear fluorescence of a single cell (n = 3 independent experiments). **(G)** Strand- and segment-specific RTqPCR strategy. Strand-specific reverse transcription (RT) was performed with tagged-primers to amplify separately the genomic viral RNA (vRNA) and the complementary viral RNA (cRNA). Each of the 8 IAV segments were then amplified by qPCR using the indicated specific primers. **(H)** shRNA**-**transduced and IAV-infected A549 cells (MOI 0.5) were subjected to nucleo-cytoplasmic fractionation at 6 hpi, and RNAs were extracted. vRNAs were reverse transcribed by strand-specific RTqPCR, and for each segment, the cytoplasmic and nuclear abundances were expressed as percentages of their sum. Technical triplicates from a representative experiment are shown. **(I)** Percentage of vRNAs exported to the cytoplasm (all segments pooled), from two independent experiments. Each dot represents the mean of three technical replicates per segment. **(J)** Same data as (I), shown as mean +/− range and indicating the standard deviation for each condition. In all experiments, dots represent biological replicates, each corresponding to the mean of technical triplicates. Two-tailed unpaired t tests (Student’s t-test; panels B, D, F, or Kolmogorov-Smirnov test; panels I, J) were performed using Prism 10. *P < 0.05, **P < 0.01, ***P < 0.001, ****P < 0.0001, ns: non-significant.

Being an NPC component, we hypothesised that RANBP2 may control the replication of vRNA by regulating the import of the viral polymerase after translation. Therefore, we tested the impact of RANBP2 on the nuclear import of newly synthesised PB1, PB2, and PA subunits. To focus specifically on import, we inhibited vRNP export using a pharmacological inhibitor of CRM1, KPT-330/Selinexor. First, we confirmed that KPT-330 inhibits vRNP export **(Figure S7A)**, with negligible toxicity **(Figure S7B)** and that inhibition operates from 5 hpi onwards **(Figure S7C)**. Then we performed subcellular fractionation followed by western blotting at 6 hpi to determine the localisation of the different polymerase subunits. Results revealed an increase in PB1, PB2 and PA protein levels in the nucleus **(Figure 2C-D)**, confirmed by the immunofluorescence detection of nuclear PB1 **(Figure 2E-F)**, indicating that the import of the neo-synthesised polymerase is facilitated in the absence of RANBP2.

### RANBP2 depletion favours the export of vRNA segments to the cytoplasm and disrupts the segment stoichiometry

Having determined that RANBP2 controls IAV replication in the nucleus by regulating the import of the newly synthesised viral polymerase, we asked why the increase in vRNA and vRNP does not translate into an increased production of infectious particles **(Figure 1A)**. To address this, we performed strand-specific RT to distinguish vRNA and cRNA, as previously **(Figure S2A)**, then quantified the 8 IAV segments by qPCR to investigate segment stoichiometry **(Figure 2G, Figure S8A-B)**. After sub-cellular fractionation, analysis of viral RNA localization revealed that, while RANBP2 knockdown led to an overall increase of all cRNA segments in the nucleus **(Figure S9)**, it also led to an overall increase in the nuclear export of vRNA segments **(Figure 2H-I)**. This effect disproportionally affected individual segments across experiments, as reflected by a higher standard deviation **(Figure 2J)**. These findings indicate that, while the depletion of RANBP2 increases the replication of all vRNA segments, it also promotes their uneven export to the cytoplasm, thereby disrupting segment stoichiometry.

Taken together, our results indicated that the knockdown of RANBP2 perturbs infection by IAV, first by facilitating the re-import of polymerase into the nucleus, which increases cRNA and vRNA, second by disproportionately favouring the export of vRNA segments into the cytoplasm. The dysregulation of both these steps may cause an abnormal accumulation of some vRNA segments in the cytoplasm, thus constituting potential pathogen-associated molecular patterns (PAMPs) that may be sensed by the infected cell.

### RANBP2 knockdown exacerbates the inflammatory response to IAV infection in primary human macrophages

To investigate whether the abnormal cytoplasmic accumulation of vRNA, observed in RANBP2-depleted cells, might associate with increased inflammation, we measured the expression of inflammatory transcripts by qPCR. In both A549 and U2OS cells, RANBP2 depletion led to an upregulation of inflammatory transcripts, which was significant in both cell types for IL-6 and IL-1β **(Figure 3A-B, Figure S10A-B)**. These changes mirrored the increase in vRNA (**Figure S10C**) and were confirmed using an independent RANBP2-targeting shRNA **(Figure S11)**. Interestingly, a modest increase in some of these transcripts was also observed in uninfected cells **(Figure 3A-B)**, likely reflecting low level activation triggered by lentiviral vector transduction, but this was not significant.

**Figure 3:**
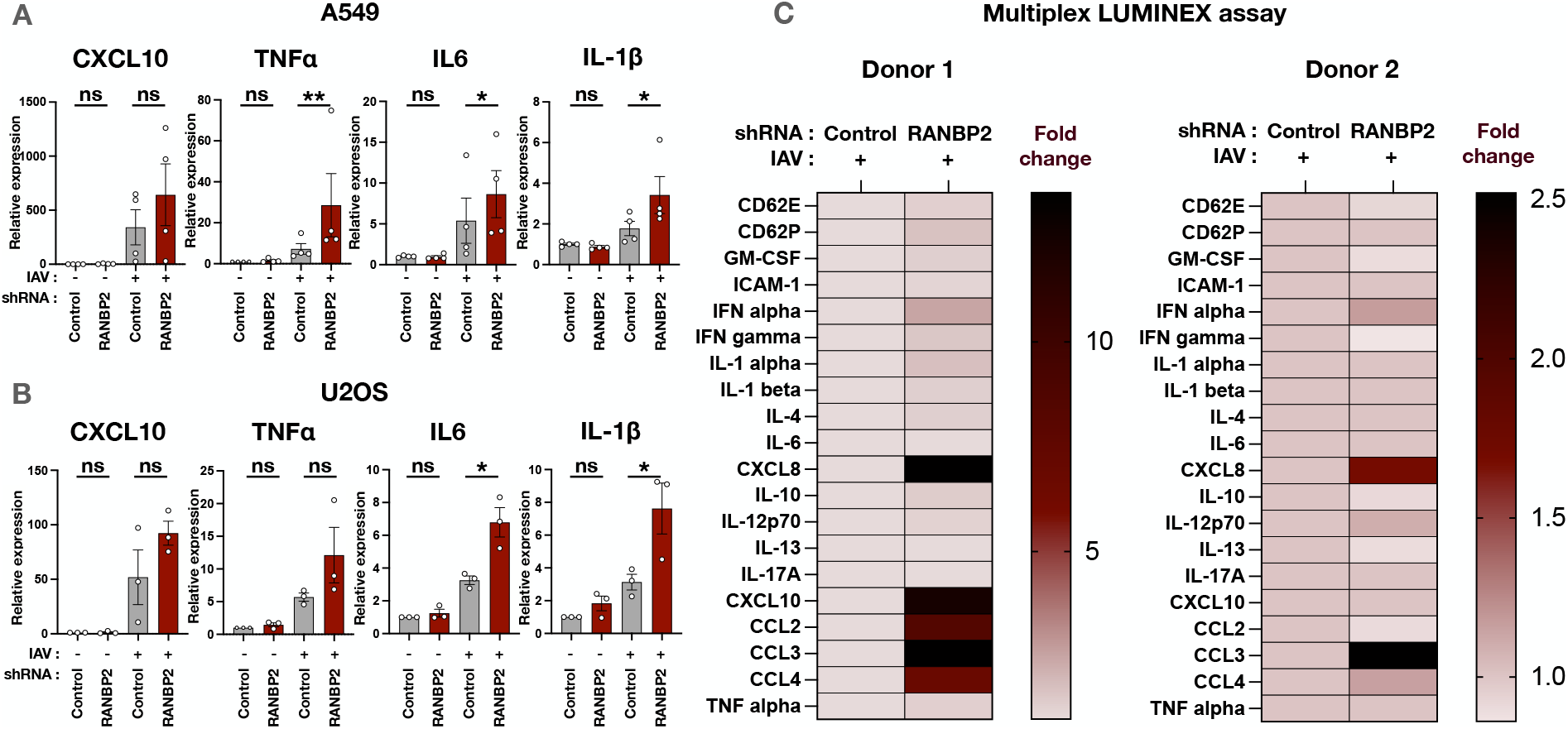
RANBP2 knockdown exacerbates the inflammatory response to IAV infection. **(A)** A549 and **(B)** U2OS cells transduced with LV shRNA-control or shRNA-RANBP2 (MOI 30) were infected with IAV (A/WSN/1933; MOI 0.5) overnight. Cytokine transcripts were quantified by RT-qPCR and results were normalized on RPL13a and to the unstimulated control condition. **(C)** Supernatants from IAV-stimulated monocyte-derived macrophages (MDM) from two donors were collected at 24 h post-stimulation and pro-inflammatory mediators were quantified by multiplex Luminex assay on samples diluted 1:50. The mean fluorescence intensities (MFI) for RANBP2-KD samples were normalized on the control samples. Results are represented as heat maps of fold values. In qPCR quantifications (panels A, B), dots represent independent biological replicates, each corresponding to the mean of technical triplicates normalized to a housekeeping gene (ΔCt). Data are plotted relative to the control unstimulated condition (2^-ΔΔCt) and shown as mean +/− SEM. Statistical analyses were performed on biological replicates only. Two-tailed paired Student’s t tests were performed using Prism 10. *P < 0.05, **P < 0.01, ns: non-significant.

To confirm this at the protein level, primary monocytes were isolated from the PBMCs of healthy donors and differentiated into pro-inflammatory M1-like macrophages. After 7 days of differentiation, primary monocyte-derived macrophages (MDM) were CD3-CD14+ CD16+ CD11b+ HLA-DR+ CD80- **(Figure S12)**. We first determined that IAV can infect MDMs **(Figure S13A)**, and stimulate an effective immune response without additional pre-stimulation, triggering the synthesis of pro-inflammatory cytokines such as IL-6 and IL-1β **(Figure S13B)**. MDM were then transduced with LVs coding for control or RANBP2 shRNA, and infected with IAV **(Figure S14A-B)**. Supernatants were collected at 24 hpi and the secretion of pro-inflammatory mediators was analysed by Multiplex Luminex assay using the manufacturer’s pre-defined inflammatory panel. Strikingly, the knockdown of RANBP2 in the primary macrophages from two donors led to an exacerbated inflammatory response to IAV infection, with a stronger induction of the pro-inflammatory chemokines CXCL8, CXCL10, CCL2, CCL3 and CCL4 **(Figure 3C)**.

Together, these findings suggest that RANBP2 plays a critical role in modulating the inflammatory response to IAV infection by regulating vRNA replication, nucleocytoplasmic trafficking of viral polymerase subunits, and selective vRNA export. Results suggest that the abnormal accumulation of viral components in the cytoplasm amplifies inflammatory signalling, highlighting a potential mechanism underlying the pathogenesis of ANE.

### The RANBP2-T585M ANE1 variant drives IAV replication and hyperinflammation following infection

Having identified a role of RANBP2 in controlling the IAV-triggered inflammation, we investigated if this function is compromised by the predominant ANE-associated mutation, c.C1754T, p.T585M. Heterozygous mutations in *RANBP2* are associated with increased susceptibility to ANE, however, it is not known how these affect protein function. Indeed, previous work showed that ANE1 mutations do not alter the structure of RANBP2 (Bley et al., 2022), and that exogenously-expressed RANBP2-T585M still localises to the nuclear envelope (Bley et al., 2022; Shen et al., 2021).

We introduced the predominant mutation (c.C1754T, p.T585M) by CRISPR-Cas9 knock-in of U2OS cells **(Figure 4A)**. A total of 47 clones were isolated after puromycin selection during 3 weeks. Mutations were confirmed by allelic qPCR **(Figure 4B, Figure S15A-C)**, using a previously published protocol (Gouy et al., 2023), and by Sanger sequencing of the cDNA **(Figure 4C-D)**. For many clones, we noted that CRISPR repair had occurred using the highly homologous RGPD sequences present on the same chromosome (Desgraupes et al., 2023), rather than the repair plasmid, therefore only clones that did not recombine with RGPD were selected **(Figure S15B)**. In total, two wild-type clones (C4 and C15, referred to as WT/WT), one heterozygously-mutated (C10, WT/C1754T) and three homozygously-mutated (C6, C9 and C14, C1754T/ C1754T) clones were isolated.

**Figure 4:**
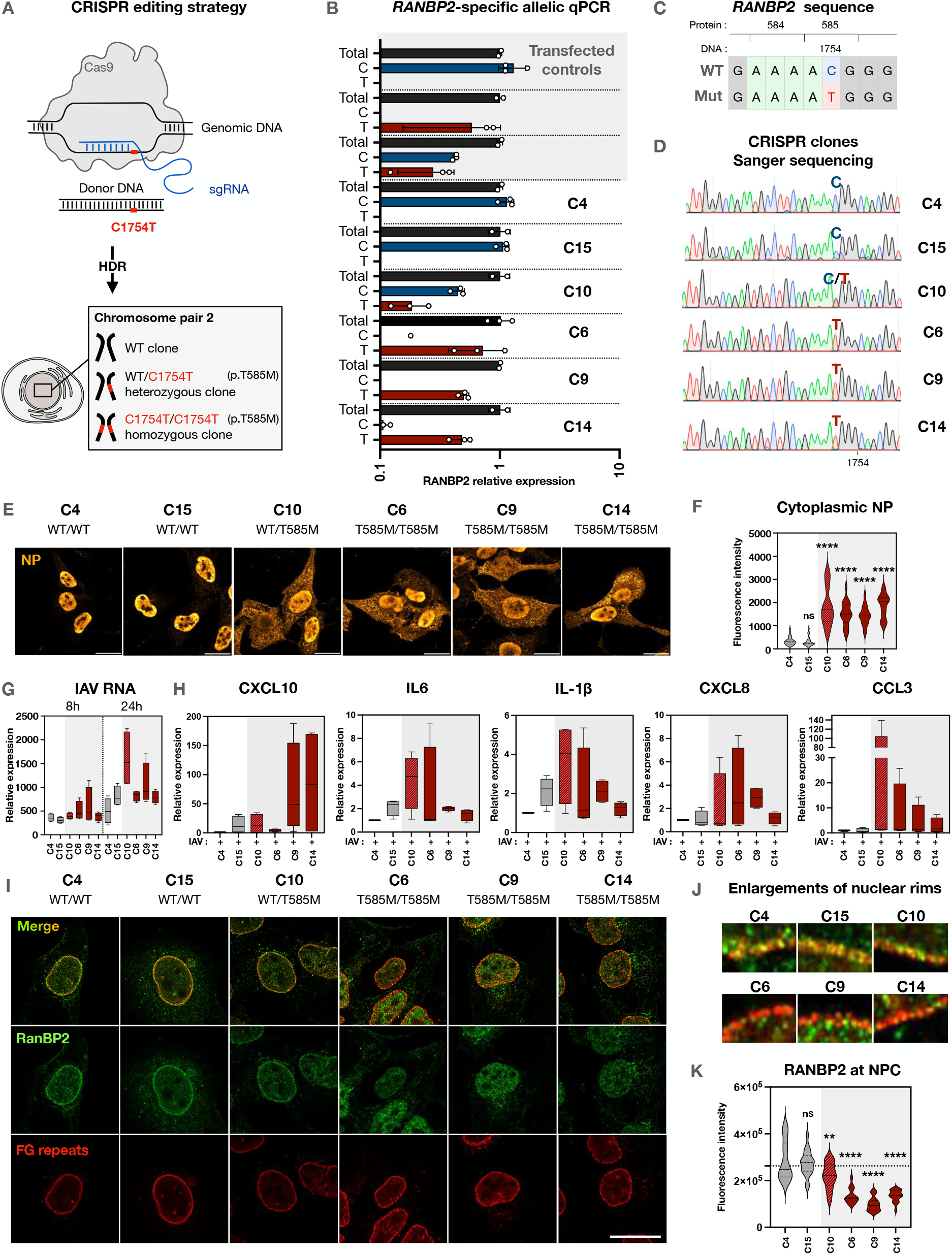
Mislocalisation of the RANBP2-T585M ANE1 variant amplifies IAV replication and hyper-inflammation. **(A)** Schematic depiction of the CRISPR editing strategy. A guide sgRNA hybridizing directly upstream the C1754 position was used to induce the cleavage by the Cas9 enzyme. A donor DNA sequence carrying the C1754T mutation (corresponding to T585M at the protein level) was supplied. After homology directed repair (HDR), three types of CRISPR clones were generated (box below): (top) WT homozygous clones (middle) WT/C1754T heterozygous clones and (bottom) C1754T/C1754T homozygous clones. **(B)** RNA expression of the WT or C1754T form of RANBP2 was assessed by RANBP2-specific RTqPCR. HEK-293T cells transfected with plasmids encoding the WT form, the T585M form or both forms of RANBP2 were used as control. **(C)** Illustration of the *RANBP2* sequence from positions 1749 to 1757, carrying a C on the WT sequence at DNA position 1754, replaced by a T on the mutant. This mutation on DNA (c.C1754T) leads to the T585M mutation on the RANBP2 protein (p.T585M). **(D)** Sanger sequencing of the CRISPR clones showing either a C, a T or both at position 1754. **(E)** CRISPR clones were infected with IAV (A/WSN/1933, MOI 0.5) and stained for NP at 8 hpi. The images from 6 additional representative fields are provided for each clone in Figure S16. **(F)** Quantification of cytoplasmic NP fluorescence. **(G)** CRISPR clones were infected with IAV (A/WSN/1933, MOI 0.5). At 1, 8 and 24 hpi, total intracellular RNAs were extracted and viral RNAs encoding the Matrix 1 (M1) protein were amplified by RTqPCR. For each clone, results were normalized on the corresponding early time point (1h). Statistical significance by two-way ANOVA is provided in Figure S17. **(H)** CRISPR clones were infected overnight with IAV (A/WSN/1933, MOI 0.5) and total intracellular RNAs were extracted. Pro-inflammatory cytokine expression was assessed by RTqPCR to detect CXCL10, IL-6, IL-1β, CXCL8 and CCL3 transcripts. For qPCR quantifications (panels G, H), results show the mean of n = 4 independent experiments, each corresponding to the mean of technical triplicates normalized to RPL13a (ΔCt). Data are plotted relative to the unstimulated condition (2^-ΔΔCt) and normalized on the WT clone C4. Results are shown as mean +/− SEM. Statistical significance by one-way ANOVA is provided in Figure S18. **(I)** CRISPR clones were fixed and stained for RANBP2 (Proteintech 27606-1-AP) and FG-repeats (Biolegend mab414). **(J)** Enlarged views of nuclear rims from panel I. **(K)** RANBP2 fluorescence was quantified at the nuclear rim (defined using the mab414 staining). Confocal images (panels E, I, J) are representative of n = 3 independent experiments. All scale bars: 20µm. Imaging quantifications (panels F, K) were performed using Fiji and are shown as violin plots (n = 3 independent experiments). One-way ANOVA (panels F, K) was performed using Prism 10. **P < 0.01, ****P < 0.0001, ns: non-significant.

CRISPR-knock-in clones were infected with IAV, and replication was assessed by quantifying viral NP at 8 hpi, as previously **(Figure 1G-H)**. Strikingly, cytoplasmic NP was significantly increased in all ANE1 CRISPR clones compared with wild-type clones, regardless of whether the mutation was heterozygous or homozygous **(Figure 4E-F, Figure S16)**. qPCR analysis confirmed that both heterozygous and homozygous ANE1 genotypes exhibited a greater increase in viral RNA replication over time, with significantly higher vRNA levels at 24 hpi compared to WT clones **(Figure 4G, Figure S17)**. Moreover, IAV infection induced an overall upregulation of inflammatory cytokines and chemokines, including CXCL10, IL-6, IL-1β, CXCL8 and CCL3, with significantly higher overall expression in both heterozygous and homozygous ANE1 genotypes compared to WT clones **(Figure 4H, Figure S18)**. Together, these results indicate that the ANE1-associated T585M mutation phenocopies RANBP2 knockdown by enhancing IAV replication and associated hyperinflammation.

### In CRISPR-Cas9 T585M knock-in cells, RANBP2 mislocalises away from nuclear pores

Although previous studies reported that exogenously-expressed RANBP2-T585M localizes to the NPC (Bley et al., 2022; Shen et al., 2021), its expression and localization have not been examined in genetically edited cell lines. Overall, we did not observe a statistically significant variation in the protein levels of RANBP2 **(Figure S19A-B)**. We next examined the subcellular localisation of RANBP2 by immunofluorescence in the clones expressing RANBP2-T585M heterozygously and homozygously using a RANBP2-specific antibody recognising the CTD. As expected, RANBP2 localised to the nuclear envelope in WT clones. In contrast, all three clones homozygous for RANBP2-T585M lost the characteristic nuclear rim staining, indicating that the mutation disrupts RANBP2 retention at the nuclear envelope **(Figure 4I-4J)**. Despite this, nuclear pores appeared intact, since the labelling of FG-repeat Nups with the MAb414 antibody was comparable across all clones **(Figure 4I)**. In heterozygous clones, RANBP2 did localise at the nuclear envelope, although quantification revealed a modest but significant decrease compared to control clones, suggesting an intermediary phenotype **(Figure 4K)**.

In conclusion, our findings suggest that RANBP2 localization at the nuclear envelope is critical to safeguard cells from the pathological inflammation following viral infection. In ANE1, mutations in the NTD of RANBP2 are specifically associated with hyperinflammation.

## DISCUSSION

In this work, we show that the NPC component RANBP2 regulates the nucleocytoplasmic transport of IAV proteins and RNAs. Its absence from nuclear pores leads to exacerbated viral replication in the nucleus and dysregulated export of vRNA to the cytoplasm, resulting in the abnormal accumulation of vRNA segments that fail to support productive particle release. While previous work indicated that RANBP2 is a co-factor for the infection by some viruses (Bichel et al., 2013; Di Nunzio et al., 2012; Schaller et al., 2011; Zhang et al., 2010; Copeland et al., 2009; Hofemeister and O’Hare, 2008), this study reveals an opposite role in IAV infection, namely the restriction of key nuclear transport steps.

RANBP2 can regulate the nucleocytoplasmic transport of macromolecules by two major mechanisms, first by regulating the cycle of Ran, second by interacting with nuclear transport receptors (Yokoyama et al., 1995; Hutten et al., 2008). The replication of IAV involves four distinct sequential NCT steps, namely the import of incoming vRNPs, the export of viral mRNAs, the import of translated viral proteins, and the export of vRNPs. RANBP2 was found to regulate only two of these steps, suggesting a specific mechanism. Specifically, RANBP2 was not involved in the initial entry of incoming vRNPs into the nucleus, which involves the classical karyopherin α/β nuclear import pathway (Miyake et al., 2019; Nguyen et al., 2023; Dou et al., 2018), and did not seem to affect the initial export of viral transcripts, which involves the general mRNA export pathway via NXF1:NXT1 (Bhat et al., 2023; Pereira et al., 2017; Zhang et al., 2024; Zhao et al., 2022). Rather, RANBP2 acted on late nuclear transport steps of IAV, after translation of the viral proteins. First, RANBP2 was found to regulate the re-import of the polymerase complex subunits, which is known to involve different pathways, the best characterised being via the β-karyopherin RanBP5 (Huet et al., 2010; Hemerka et al., 2009; Cros et al., 2005). Second, results suggest that RANBP2 exerts a quality control checkpoint for vRNAs that are exported back to the cytoplasm, a step known to depend on the β-karyopherin CRM1 (Elton et al., 2001; Ma et al., 2001; Watanabe et al., 2001). Our results therefore suggest that RANBP2 may impact viral NCT by interacting with key β-karyopherins.

Although vRNA levels increased in RANBP2-deficient cells, infectious viral particle production did not. This likely results from the disproportionate accumulation of some vRNA segments in the cytoplasm observed in RANBP2-KD cells, which could impair packaging. The resulting cytoplasmic vRNA can activate RIG-I, which results in the production of pro-inflammatory cytokines through NF-κB signalling (Iwasaki and Pillai, 2014). Consistent with this, RANBP2-KD cells exhibited a pronounced inflammatory response following IAV infection, particularly marked by elevated chemokine production and secretion. These findings suggest that the abnormal cytoplasmic accumulation of vRNA segments may act as PAMPs that amplify the inflammatory response during IAV infection.

Influenza viruses are the predominant viral trigger of ANE1 and reasons for this remain unclear (Bartolini et al., 2024; Jiang et al., 2022; Mizuguchi et al., 2007; Neilson et al., 2003; Chatur et al., 2022; Bashiri et al., 2020). In particular, the high prevalence for Influenza over other respiratory infections (e.g., RSV, SARS-CoV-2) and other childhood febrile illnesses (e.g., streptococcal infections) is puzzling and unexplained. Our work suggests that, by restraining key steps of IAV NCT, RANBP2 is required to limit the accumulation of viral PAMPs in the cytoplasm.

*RANBP2* pathogenic variants are known to be heterozygous dominant point mutations, yet their underlying molecular mechanism remains not known. We previously reported that both wild-type and mutant *RANBP2* alleles are co-expressed in ANE1 patients (Gouy et al., 2023), suggesting a possible gain-of-function or dominant-negative effect. However, exogenous T585M RANBP2 is expressed normally and localises correctly to the nuclear envelope (Bley et al., 2022; Shen et al., 2021), arguing against a dominant-negative mechanism. Still, overexpression studies may not reflect the physiological balance of wild-type and mutant protein in patients, and the use of artificial tags could perturb natural functions and interactions.

Here, using CRISPR knock-in clones, we demonstrate that the T585M mutation prevents RANBP2 from properly associating with the nuclear envelope, which is consistent with the fact that most ANE1 variants cluster within the NTD responsible for NPC anchoring. Importantly, the T585M mutation phenocopied RANBP2 knockdown in their effects of viral replication and inflammatory responses, supporting a haploinsufficiency mechanism that reflects a specific loss of RANBP2 function at the nuclear envelope rather than a global reduction in RANBP2 protein. In heterozygous cells, only partial RANBP2 localisation to the nuclear envelope was observed, further supporting this model. Notably, RANBP2 has a pLI (probability of Loss-of-function Intolerance) score of 1 in gnomAD, indicating strong intolerance to loss of function variants, and confirming the essential role of its proper expression and localisation at the nuclear pore complex in ANE1 pathogenesis.

This study provides the first direct evidence that an ANE1-associated mutation enhances IAV-induced inflammatory responses. This phenotype was equally pronounced in homozygous and heterozygous ANE1 clones, indicating that even partial mislocalisation of RANBP2 can markedly amplify IAV-triggered inflammation. This role of RANBP2 in restraining virus-induced hyperinflammation is consistent with the elevated pro-inflammatory cytokine levels reported in ANE1 patients, most of whom carry heterozygous RANBP2 mutations, with only a single case of biallelic case described (Ayaz et al., 2022). However, this work does not yet explain how increased peripheral inflammation leads to encephalopathy in patients. Future studies in individuals carrying ANE1 mutations will be essential to define how these variants affect the composition and function of innate immune cell populations. Moreover, since there is currently no evidence of innate immune cell infiltration in patient brains during ANE1 episodes, animal models will be critical to assess how neurons and glia contribute to disease onset.

## MATERIALS & METHODS

### Viruses, lentiviral vectors and cells

Infections with IAV were carried out with the H1N1/A/WSN/1933 strain in all experiments, except for the Alpha-Centauri assay, where this same strain was engineered to express a fragment of the NanoLuc (IAV-α), as described below. Lentiviral vectors (LV) were produced in HEK-293T cells (ATCC) by calcium phosphate transfection using a VSV-G envelope plasmid (pVSVg; 15 µg in a 70-80% confluent 175cm^2^ flask), a Gag-Pol encapsidation plasmid (p8.74; 30 µg), and the transfer plasmid of interest (30 µg; mCherry-shRNA-control, mCherry-shRNA-RANBP2 or Cen-NLS; see **Table S1**). After 48 h, vectors were harvested and ultracentrifuged for 1 h at 22,000 rpm (Optima XE-90, Beckman Coulter) at 4°C. A549 cells (ATCC) and CRISPR-engineered U2OS clones (generated in-house) were cultured in Dulbecco’s Modified Eagle Medium (DMEM, Gibco) supplemented with 10% of Fetal bovine serum (FBS, Serana) and 1× penicillin/streptomycin (PS, Gibco) at 37°C with 5% CO2. THP-1 cells and monocyte-derived macrophages (MDM) were cultured in Roswell Park Memorial Institute (RPMI-1640, Gibco) medium supplemented with 10% FBS and 1× of PS. MDCK cells (ATCC) were cultured in Minimum Essential Medium (MEM, Gibco) supplemented with 5% FBS and 1× of PS.

### Lentiviral transduction

Cells were seeded into plates one day prior to transduction. After a PBS wash, lentiviral transduction was performed at MOI 20 to 40 (transducing units/cell) in a small volume of culture medium supplemented with 2% FBS, to facilitate viral adsorption. After 2 h of incubation at 37°C, 10% FBS culture medium was added. Transduced cells were cultured for 48-72 h prior to viral infections to allow efficient knockdown. All experiments were completed by 56-96 h post-transduction.

### Viral production

H1N1/A/WSN/1933 viral stocks were produced by amplification at an MOI of 10^−4^ on MDCK cells, cultured in MEM containing 2%-FBS and 1 μg/mL of TPCK-treated trypsin for 3 days at 37°C. Viral supernatants were harvested and centrifuged for 5 min at 2,000 g to remove cellular debris. Aliquots were stored at −80°C. IAV titers were determined by TCID50 assay as described below.

### IAV infections

Cells were infected with IAV at MOI 0.5 for all experiments, except when monitoring the early steps of the viral cycle (i.e. internalization and nuclear import), where MOI 4 was used to increase signal detection. Cells were exposed to IAV for 1 h in a small volume of 2% FBS culture medium to promote viral adsorption at the cell surface before replacing the inoculum with warm culture medium and incubating at 37°C with 5% CO2.

### RNA extractions and RTqPCR

Intracellular RNA was extracted using the RNeasy mini kit (Qiagen) according to the manufacturer’s protocol. RNA yield and purity were assessed by spectrophotometry (NanoDrop 2000c, Thermofisher Scientific). Reverse transcription (RT) using oligo(dT) and random primers was performed using the PrimeScript RT Reagent Kit (Perfect Real Time, Takara Bio Inc). Strand-specific RT for IAV RNA was performed using tagged primers that bind either to the consensus 3’-end sequence of vRNAs or cRNAs of all 8 segments, with Superscript III (Invitrogen, Thermofisher Scientific) in the presence of 0.1M DTT (Invitrogen, Thermofisher Scientific), 10mM dNTP (Invitrogen, Thermofisher Scientific) and RNAsin (Promega). Segment distinction was then achieved by qPCR using specific primer pairs. RANBP2-specific RT was performed using a primer (1µM) binding to a region absent from RGPDs, as described in (Gouy et al., 2023) and the PrimeScript RT Reagent Kit. Real-time quantitative PCR was performed using the Power Up kit (Applied Biosystems, Thermofisher Scientific) on the ViiA7 thermocycler (Life technologies, Thermofisher Scientific). All RT and qPCR primers are listed in **Table S2**.

### Immunofluorescence

Cells were washed with PBS and fixed with 4% paraformaldehyde (PFA, Thermofisher Scientific) for 10 min at room temperature (R/T). After 3 PBS washes, remaining PFA was quenched with PBS NH4Cl 50 mM for 10 min at R/T. Cells were then permeabilized using PBS 0.5% Triton-100X (Sigma Aldrich) for 15 min at R/T, and saturated in PBS 2% Normal goat serum (NGS, Invitrogen, Thermofisher Scientific) with 2 PBS washes between each step. Incubation with primary antibodies for 1 h at R/T was followed by 5 PBS washes and incubation with secondary antibodies for 30 min at R/T (**Table S3**). After further washing, cells were stained with Hoechst (1:10,000 dilution in PBS; Thermofisher Scientific) for 5 min at R/T, washed, and mounted in Prolong Diamond Antifade mounting medium (Invitrogen, Thermofisher Scientific). Images were acquired on a Zeiss LSM880 Airyscan confocal microscope at the Montpellier Ressources Imagerie (MRI) facility and processed using Fiji (2.14.0). For visualisation, Alexafluor-488 signal corresponding to NP was pseudocoloured (“Orange Hot”) and brightness was increased uniformly across conditions.

### Alpha-Centauri assay

Cloning of IAV-α. A DNA fragment coding for αNluc (GVTGWRLCERILA) flanked by NotI and NheI overhangs was obtained by annealing two overlapping oligonucleotides:

**NotI-αNLuc-NheI-F** GGCCGCAGGGGGAGTGACAGGGTGGAGACTATGCGAAAGAATACTTGCATAAG

**NotI-αNLuc-NheI-R** CTAGCTTATGCAAGTATTCTTTCGCATAGTCTCCACCCTGTCACTCCCCCTGC

The reverse genetics plasmid pPolI-SL-PB2-αNluc was generated by subcloning this fragment between the NotI and NheI sites of the pPolI-SL-PB2-Nanoluc (Diot et al., 2016).

Particle production by reverse genetics. The eight pPolI-WSN plasmids (-PB2-αNluc, -PB1, -PA, -HA, -NP, -NA, -NS, -M), and four expression plasmids (pcDNA3.1-WSN-PB2, -PB1, -PA, -NP; 0.5 μg of each) were co-transfected into a co-culture of 293T and MDCK cells (seeded in a 6-well plate at 4 × 10^5^ and 3 × 10^5^ cells, respectively) using 10 μL of FuGENE® HD transfection reagent (Promega). After 24 h at 35°C, cells were washed twice with DMEM and incubated in DMEM containing 1 μg/mL of TPCK-treated trypsin for 48 h. Recombinant PB2-αNluc virus was amplified at an MOI of 10^−4^ on MDCK cells for 3 days at 35°C. Viral stocks were titrated on MDCK by plaque assays and sequenced to confirm the presence of the αNluc coding sequence.

Cells were transduced with LV Cen-NLS for 3 days, followed by LV shRNA-control or LV shRNA-RANBP2 at MOI 15 for 4 days. Cells were then infected with IAV-α at MOI 4 for the indicated times. Luminescence was measured using NanoGlo Live Cell substrate (Promega) and an Infinite M Plex spectrophotometer (Tecan).

### Nucleo-cytoplasmic fractionation

Nuclear and cytoplasmic fractions were separated using the NE-PER Kit (Thermofisher Scientific) according to the manufacturer’s protocol. Briefly, 1-2 × 10^6^ cells were washed in PBS, lysed in cold CER I buffer, vortexed 15 s, and incubated on ice for 10 min. Cold CER II buffer was then added, followed by brief vortexing and incubation on ice for 1 min. After 5 s vortexing and full speed centrifugation, supernatants (cytoplasmic fraction) were collected and stored at −80°C. Nuclear pellets were resuspended in cold NER buffer and subjected to 4 cycles of 15 s vortexing then 10 min incubation on ice. After full speed centrifugation, supernatants (nuclear fraction) were collected and stored at −80°C.

### Western blot

Cell lysates were sonicated for 20 cycles (30 s on/off; Bioruptor Pico, Diagenode) and mixed with 4X Laemmli buffer (250 mM Tris-HCl pH 7, 8% sodium dodecyl sulphate (SDS), 40% glycerol, 10% β-mercaptoethanol and 0.005% bromophenol blue). After denaturation at 95°C for 5 min, samples underwent SDS polyacrylamide gel electrophoresis (SureCast Gel Handcast System, Thermo Fisher Scientific) and were transferred to a 0.45 µm nitrocellulose membrane (Sigma Aldrich). Membranes were saturated in PBS containing 0.05% Tween 20 (Sigma) and 10% milk powder for 30 min, incubated with primary antibodies overnight at 4°C, washed 3 times in PBS 0.05% Tween 20, and incubated with HRP-conjugated secondary antibodies for 1 h at R/T (**Table S3**). Signals were detected with Immobilon Forte Western HRP substrate (Merck) or Immobilon ECL Ultra Western HRP Substrate (Merck) on a Chemidoc imaging system (Bio-Rad).

### TCID50 assay

MDCK cells were seeded in FBS-free MEM medium (Gibco) and inoculated with serial dilutions of viral supernatants in 2% FBS medium in octuplicates. After 3 days at 37°C and 5% CO2, cells were washed twice in PBS and fixed with 4% PFA for 10 min at R/T and stained with crystal violet. TCID50 titers were calculated as:

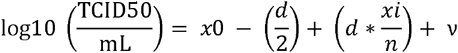

where *x*_0_ is the log of the initial dilution factor, *d* the log of the serial dilution factor, *xi* the number of positive events, *n* the number of replicates, and *ν* the log of the inoculum volume (mL).

### Peripheral blood mononuclear cell (PBMC) isolation and monocyte-derived macrophages differentiation

Buffy coats from healthy donors were obtained from the Établissement Français du Sang (EFS). Blood samples were diluted 1:1 with PBS and layered over an equivalent volume of Lymphoprep (Stemcell technologies) at R/T. Samples were centrifuged at 800 g for 30 min at 20°C (no brake). The PBMC ring was collected using a 5 mL pipet, washed 3 times in a final volume of 50 mL PBS (1,200 rpm, 5 min, 20°C with brakes on), and resuspended in 30 mL of 10% FBS RPMI-1640 medium. PBMCs were counted manually using counting chambers (Kova).

For MDM differentiation, 2 × 10^7^ cells were seeded per well into 6-well plates, allowed to adhere for 45 min, at 37°C with 5% CO2, washed 3 times with 10% FBS RPMI-1640 medium directly on the cell layers to remove loosely attached cells, and cultured in RPMI-1640 medium with 10% FBS and 50 ng/mL GM-CSF (Gentaur) for 8 days.

### Flow cytometry

After 8 days of MDM differentiation, cells were stained and analysed on a Fortessa flow cytometer (BD Biosciences) using the following antibodies: CD3-BV421 (clone UCHT1, Biolegend 300433), CD14-PerCP-Cy5.5 (clone HCD14, Biolegend 325621), CD16-Alexa700 (clone 3G8, Biolegend 302026), CD11b-APC-Cy7 (clone M1/70, Biolegend 101225), HLA-DR-FITC (REA805, Miltenyi 130-111-941), and CD80-BV650 (clone 2D10, Biolegend 305227) (see **Table S3)**.

### Multiplex Luminex assay

The presence of inflammatory mediators was assessed in supernatants of stimulated MDMs using the ProcartaPlex Human inflammation Panel 20-plex kit (Invitrogen, ThermoFisher Scientific). First, a subset of samples was centrifuged and diluted 1:10 or 1:50 to determine the optimal dilution for the assay. A 1:50 dilution was selected in order to detect the strongly induced mediators (e.g. CXCL8) within the range of the standard curves supplied in the kit. Buffers and standards were prepared according to the manufacturer’s protocol. Briefly, the Capture Bead mix was vortexed and added to the plate, followed by washing with Wash buffer, then samples and standards were added. The plate was incubated for 1 h at R/T on a shaker. After 2 washes, Biotinylated Detection Antibody mix was added for 30 min at R/T on a shaker. After 2 washes, Streptavidin-PE was added for 30 min at R/T on a shaker. After 2 more washes, the plate was read on the MAGPIX flow cytometer (Luminex).

### Establishment of U2OS clones expressing RANBP2 T585M by CRISPR knock-in

The plasmid pUcIDT Amp (synthetized by Integrated DNA Technologies) containing the donor sequence (see below) and the plasmid pSpCas9(BB)-2A-Puro (PX459; Addgene #62988) modified to contain the guide sequence (5’-GGCAGAATGCCTTCAGAAAA-3’) were co-transfected in equal amounts into human osteosarcoma U2OS cells using Lipofectamine 3000 (Invitrogen, ThermoFisher Scientific), following the manufacturer’s instructions. Two days post-transfection, cells were selected with puromycin (Gibco) at 2 µg/ml for four days. Single-cell clones were obtained by limiting dilution. A total of 198 cells were distributed in 576 wells containing 10% FBS DMEM medium. After 3 weeks, 47 clones were successfully established and analyzed by RANBP2 allele-specific RTqPCR as described above and Sanger sequencing (Eurofins).

#### Donor DNA sequence

(GCGGTTTGTACTCTGATTCACAGAAAAGCAGTgtaagtagtaaaacaaaaatattgctttcacttagtgcgtaggttttaccggggatttaatcctcatgtgaagatttaatttgtcatgtgacccattaacatatatgtatgtaagccctgaactgtgtatttagaaagcaattttagtaaattgaactattttttagACCTGGAAACGTAGCAAAATTGAGACTTCTAGTTCAGCATGAAATAAACACTCTAAGAGCCCAGGAAAAAC ATGGCCTTCAACCTGCTCTGCTTGTACATTGGGCAGA<u>g</u>TGtCT<u>g</u>CAGAAAA**t**Ggtgagttttaaagtataagcatttttaaagaa cattaccttaattttttaaaatcatgaactttttattgaaagtttttttgttctgaaaacagcagcttggtcacattatgacagatgtgttttttattgctgcaaaatagttaatgtagttaaatataagcacttagaggagcaatgcctggcacacagtgaatgttacatattagctgagctgttactgttattccttaataattaagttctgataattattcagcctgaaaattaaaaaaa)

Capital letters: exon

Lowercase: intron

Underlined: mutations

Bold lowercase: c. 1754C>T mutation

### Generation of homozygous/heterozygous RANBP2-expressing control cells for allelic qPCR

HEK-293T cells were transfected with plasmids encoding mCherry-RANBP2-shRNA and either GFP-RANBP2-WT, GFP-RANBP2-T585M, or both (**Table S1**) in calcium phosphate. After 48 h of culture, cells were detached using trypsin (Gibco) and sorted on the ARIA IIu cytometer (Becton Dickinson) at the MRI facility to isolate the mCherry-positive cells. Sorted cells were lysed and total RNAs were extracted using the RNeasy mini kit (Qiagen) according to the manufacturer’s protocol.

### MTT cell viability assay

Cell viability was assessed using the MTT [3-(4,5-dimethylthiazol-2-yl)-2,5-diphenyltetrazolium bromide] assay (Sigma). Following treatment, the culture medium was carefully removed, and cells were washes once with PBS to eliminate any residual medium or serum components. A stock solution of MTT was prepared at 5 mg/mL in DMEM without phenol red (Gibco). Subsequently, 100µl of the MTT solution was added to each well and incubated for 3 h at 37°C with 5% CO2 to allow the formation of formazan crystals by metabolically active cells. After incubation, 100µL of acidified isopropanol (1:200 HCl 0.05N) was added to each well to dissolve the formazan crystals completely. The absorbance was measured at 570 nm using an Infinite M Plex spectrophotometer (Tecan).

### Statistical analysis

Statistical analysis was performed using the GraphPad Prism software (v10.2.3). For qPCR experiments, statistical analyses were performed on ΔCt or ΔΔCt values, as indicated. ΔCt values correspond to the mean of technical replicates normalised for the housekeeping gene, and ΔΔCt values are normalised to the control condition. Unless otherwise stated, biological replicates are represented as single dots in bar graphs (mean +/− SEM). All statistical tests are specified in the figure legends.

## Supporting information

All Tables

## ACKNOWLEDGMENTS

This work was financed by grants from the Agence Nationale de la Recherche (ANR-23-CE15-0005-01) and ANRS-MIE (ECTZ209411). We thank Marion Cannac (Université de Montpellier, France) for help with phenotyping primary macrophages, and Montpellier Ressources Imagerie (MRI) for support with flow cytometry, cell sorting and imaging. We thank Jomon Joseph (National Center for Cell Science, Pune, India) for the GFP-RANBP2 construct, and Sébastien Nisole (INSERM, Montpellier, France; INRS, Institut Armand Frappier, Montréal, Canada) for the GFP-RANBP2-T585M construct.

## Supplementary Figure Legends

**Figure S1:**
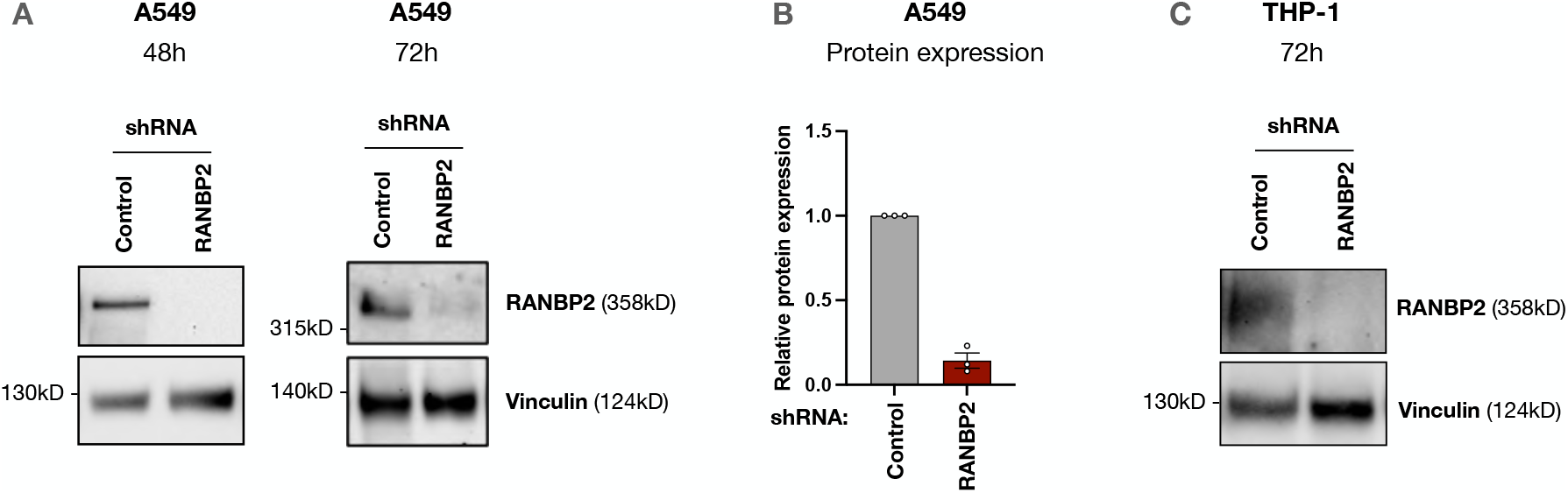
Assessment of RANBP2 knockdown in A549 and THP-1 cells. A549 and THP-1 cells were transduced with LV shRNA-control or shRNA-RANBP2 at MOI 20. **(A)** Knockdown efficacy was assessed by western blot at 48 h and 72 h post-transduction in A549 cells. **(B)** Band intensities from n = 3 independent experiments were quantified using ImageLab. Results are normalized on the vinculin signal and to the control condition. **(C)** Knockdown efficacy was assessed by western blot at 72 h post-transduction in THP-1 cells. Representative blot from n = 2 independent experiments.

**Figure S2:**
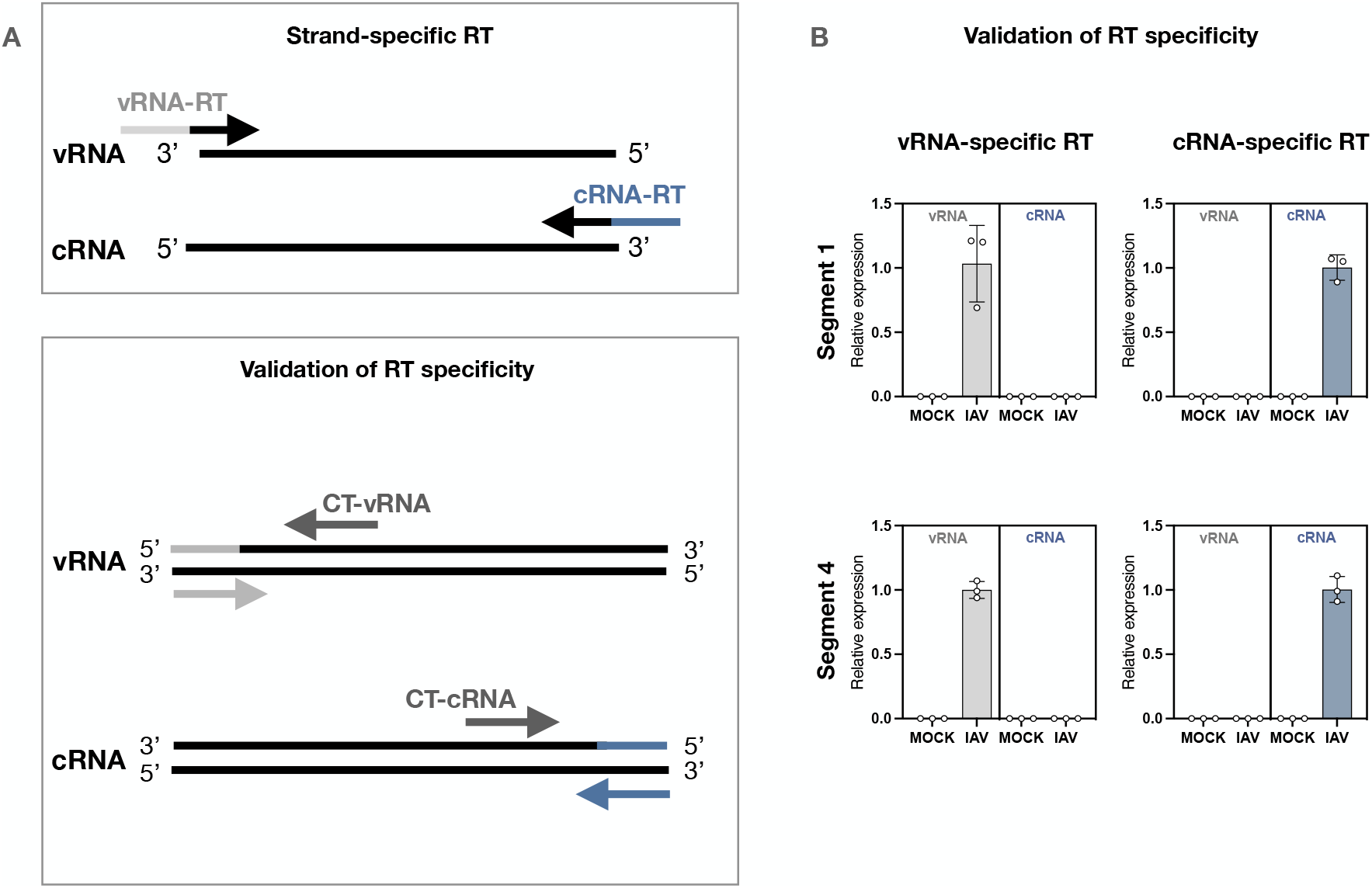
RT-qPCR strategy to distinguish cRNA and vRNA. **(A)** Strand-specific Reverse Transcription (RT) strategy. Genomic viral RNAs (vRNA) and complementary RNAs (cRNA) were reverse transcribed using specific tagged-primers hybridizing to the consensus 3’-sequences common to all 8 viral segments. **(B)** Validation of strand-specific RT specificity. qPCR primers complementary to the tags were used to amplify cDNAs produced by each RT, and to verify that only vRNAs are amplified after vRNA-specific RT, that, conversely, only cRNAs are amplified after cRNA-specific RT, and that no cellular RNAs were reverse transcribed by either RT. RT specificity was checked for two IAV segments (segments 1 and 4).

**Figure S3:**
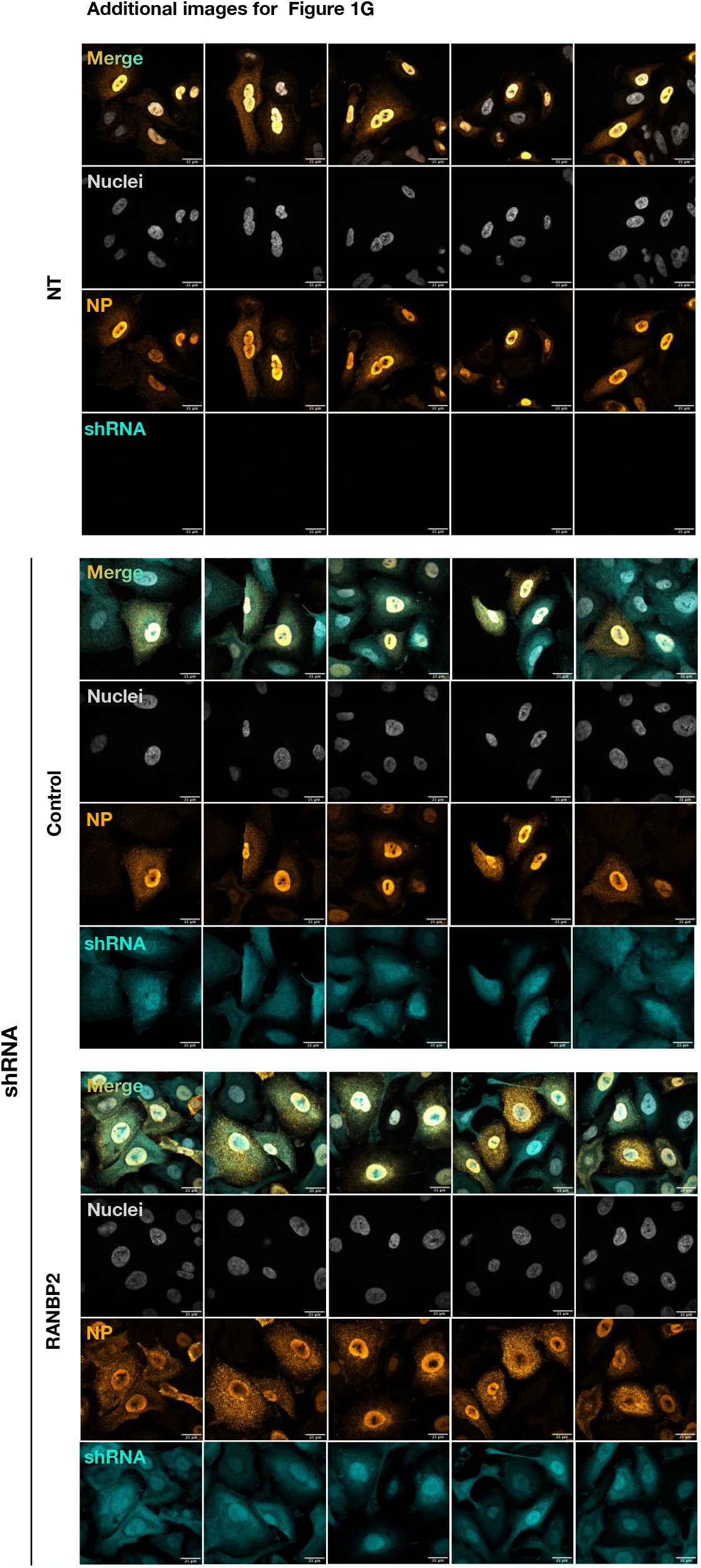
Increase in cytoplasmic NP in RANBP2-KD cells at 8 hpi. Additional representative images for Figure 1G from n = 3 independent experiments. A549 cells were transduced with LV shRNA-control or shRNA-RANBP2, or left untransduced (NT), and infected with IAV (A/WSN/1933; MOI 0.5). At 8 hpi, cells were fixed and stained for nucleoprotein (NP). Scale bar: 20µm. Image processing was performed using Fiji. NP (AlexaFluor-488) and shRNA (mCherry) signals were pseudocoloured to “Orange Hot” and “cyan” for aesthetic purposes.

**Figure S4:**
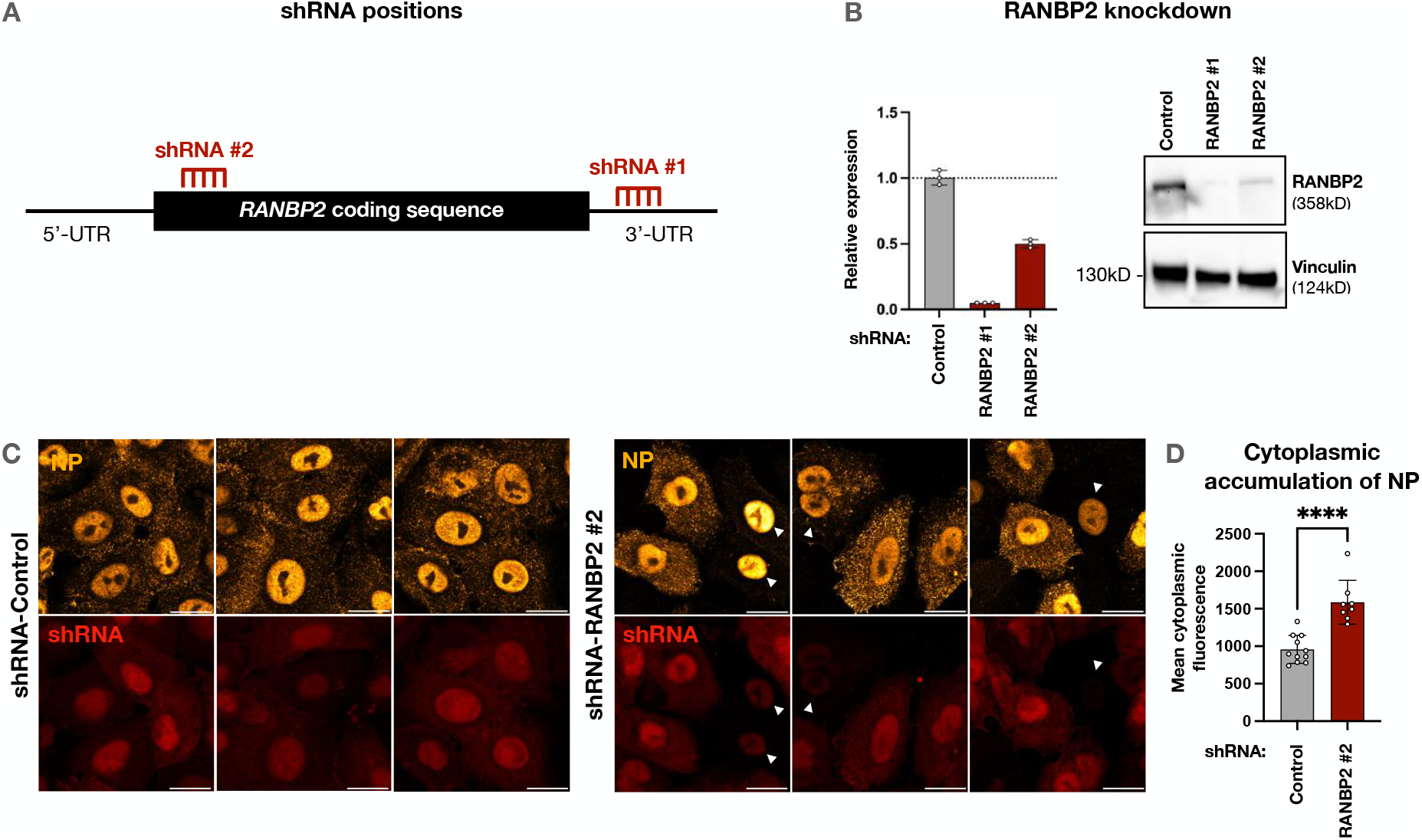
Increase in cytoplasmic NP in RANBP2-KD cells at 8 hpi, confirmed using a second shRNA. **(A)** Schematic depiction of the hybridizing sites of the two RANBP2-specific shRNAs used in this study. While shRNA-RANBP2 #1 binds in the 3’UTR sequence, shRNA-RANBP2 #2 binds at position 225 of *RANBP2* RNA. **(B)** A549 cells were transduced with LV shRNA-control or shRNA-RANBP2 for 48 h, and RANBP2 knockdown was verified by RTqPCR and western blot in n = 3 independent experiments. **(C)** Transduced A549 cells were infected with IAV (A/WSN/1933; MOI 0.5). At 8 hpi, cells were fixed and stained for NP. Scale bar: 20µm. **(D)** Cytoplasmic NP fluorescence was quantified using Fiji. Each dot represents the cytoplasmic fluorescence of a single cell. Two-tailed unpaired Student’s t test. ****P < 0.0001.

**Figure S5:**
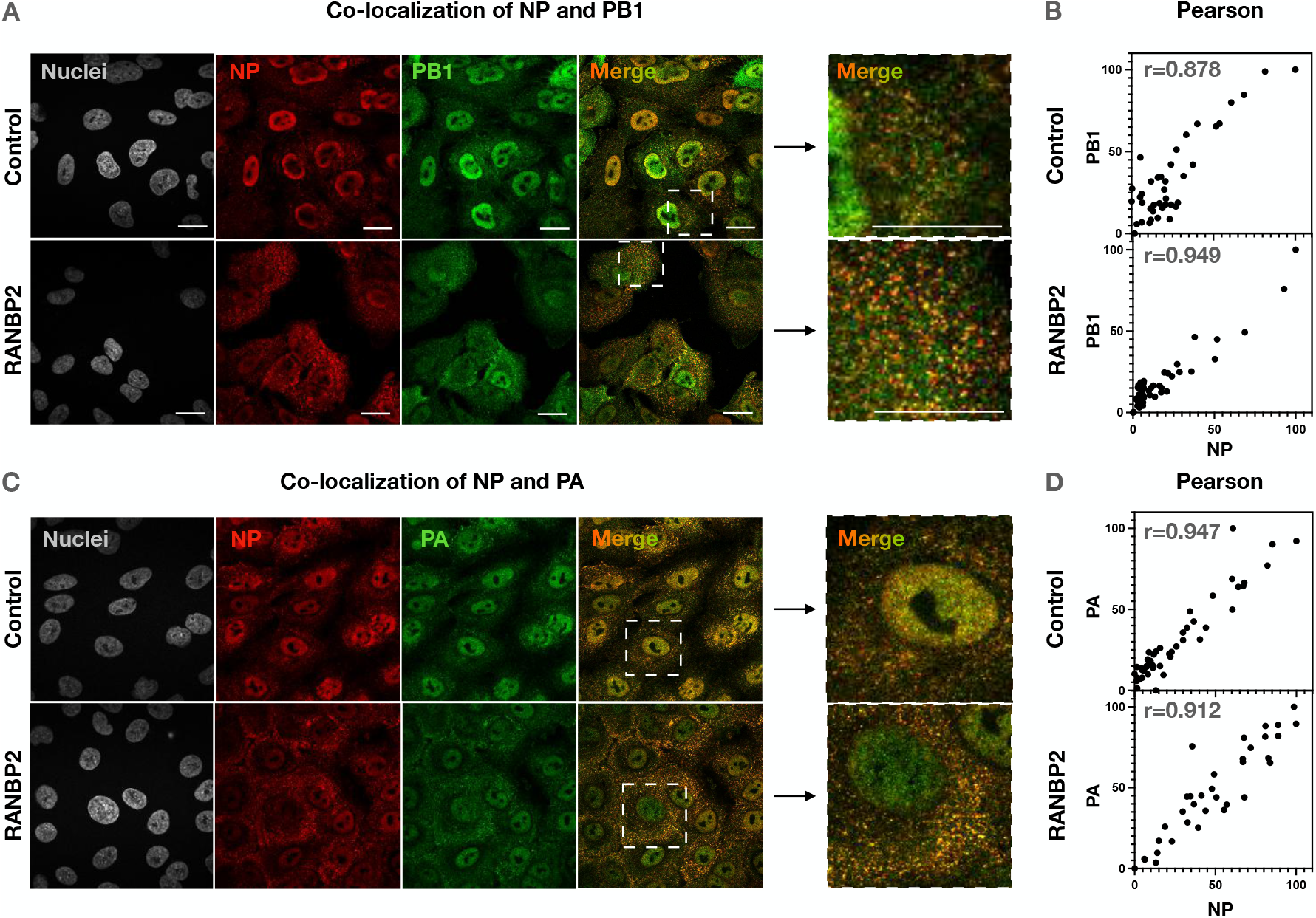
Increase in cytoplasmic vRNP in RANBP2-KD cells at 8 hpi. Transduced A549 cells were infected with IAV at MOI 0.5 and co-stained for **(A-B)** NP and PB1 or **(C-D)** NP and PA at 8 hpi. Pearson’s correlation coefficients between (B) NP and PB1 and (D) NP and PA signals were quantified using Fiji. Scale bars: 20µm.

**Figure S6:**
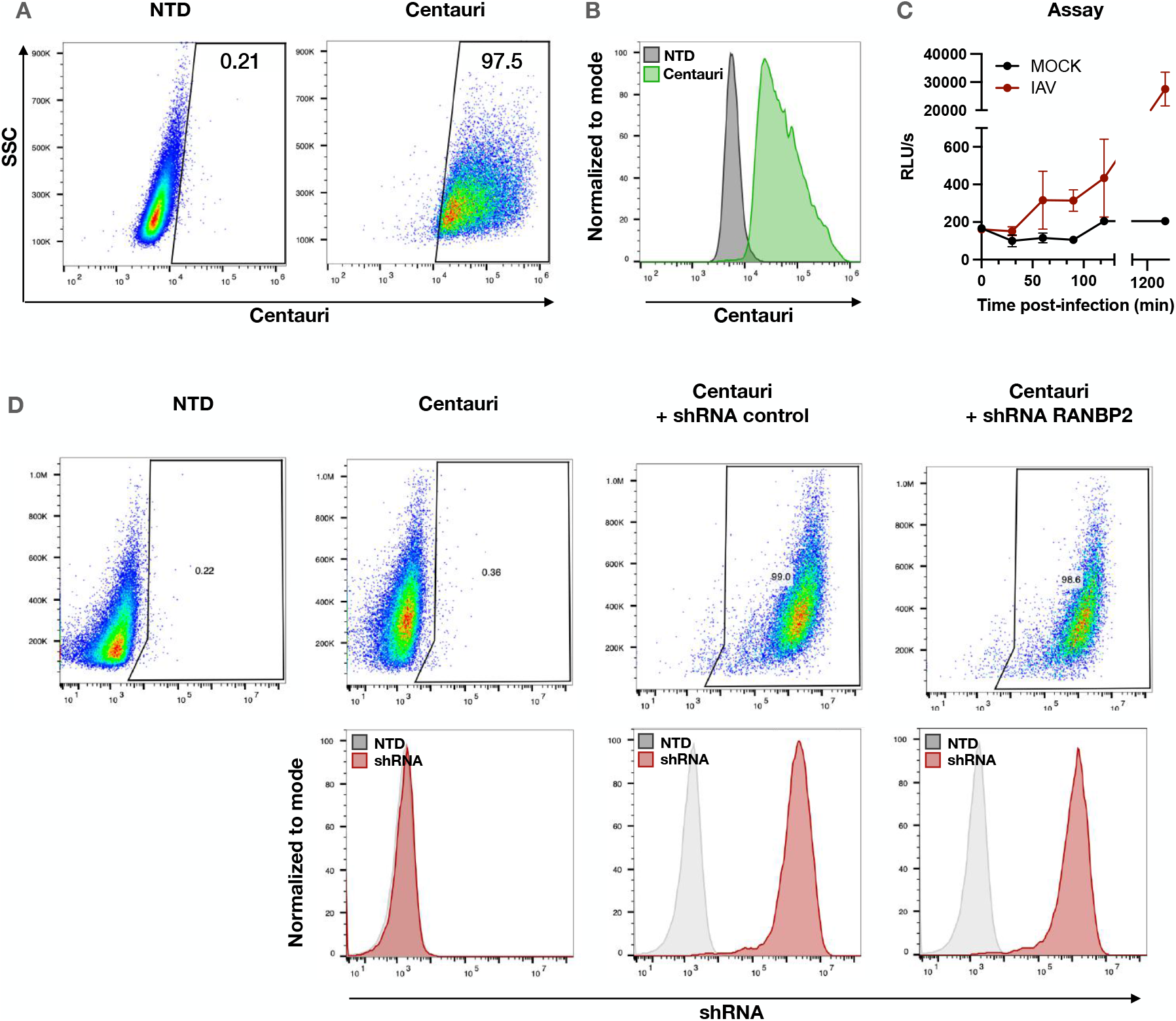
Alpha-Centauri assay to measure IAV nuclear import. **(A-B)** A549 cells were transduced with a plasmid containing the Cen-NLS tagged with HA. Cells were stained for HA and analyzed by flow cytometry to determine the percentage of transduced cells. **(C)** Cells were infected with IAV-α at MOI 4, and luminescence was measured to assess the NanoLuc complementation upon nuclear import of vRNPs. **(D)** Cen-NLS-expressing A549 cells were transduced with LV shRNA-control or shRNA-RANBP2 expressing the mCherry. The percentage of mCherry-positive cells was determined by flow cytometry.

**Figure S7:**
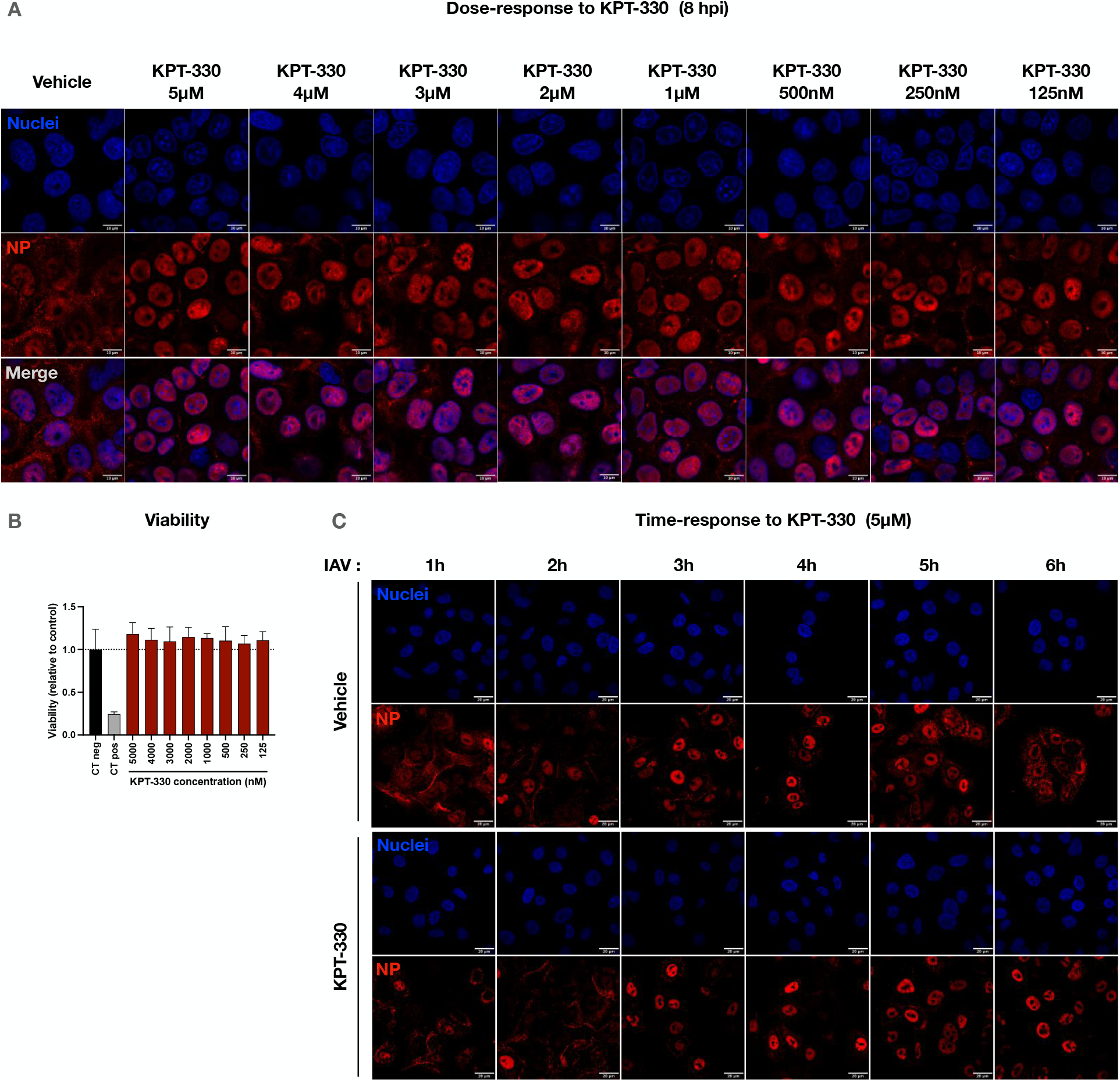
Inhibition of CRM1-dependent NP export using KPT-330/Selinexor. A549 cells were treated or not (Vehicle) with KPT-330 at the indicated doses for 2 h at 37°C, then infected with IAV. **(A)** Cells were fixed and stained for the NP at 8 hpi to verify inhibition of NP export. **(B)** KPT-330 cytotoxicity was assessed by MTT assay. Uninfected cells were incubated with MTT (5 mg/mL) for 3 h at 37°C. Acid isopropanol was added and absorbance was read at 570 nm (mean +/− SD). **(C)** At the indicated times post-infection, cells were fixed and stained for NP to determine the start of vRNP export.

**Figure S8:**
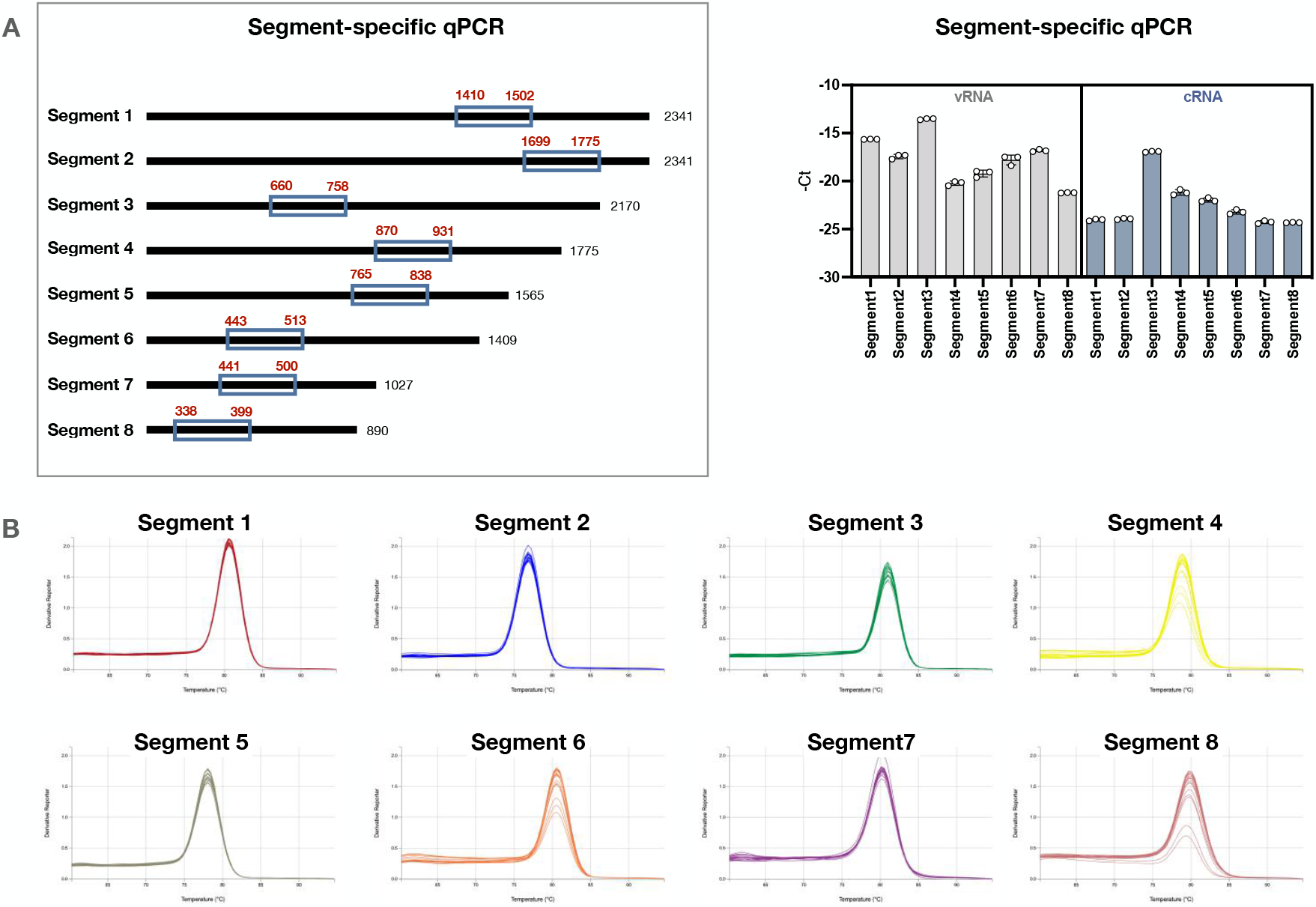
qPCR strategy to amplify the 8 segments of IAV. **(A)** Segment-specific qPCR strategy. After strand-specific RT, the 8 segments were detected using qPCR primers amplifying the indicated regions of each segment. Results from a representative experiment (technical triplicates) confirm that all 8 segments were amplified after each RT. **(B)** Melting curves indicate that all primer pairs used for segment-specific qPCR remained specific of the viral RNA segments even after total RT, which amplifies all RNAs, was performed.

**Figure S9:**
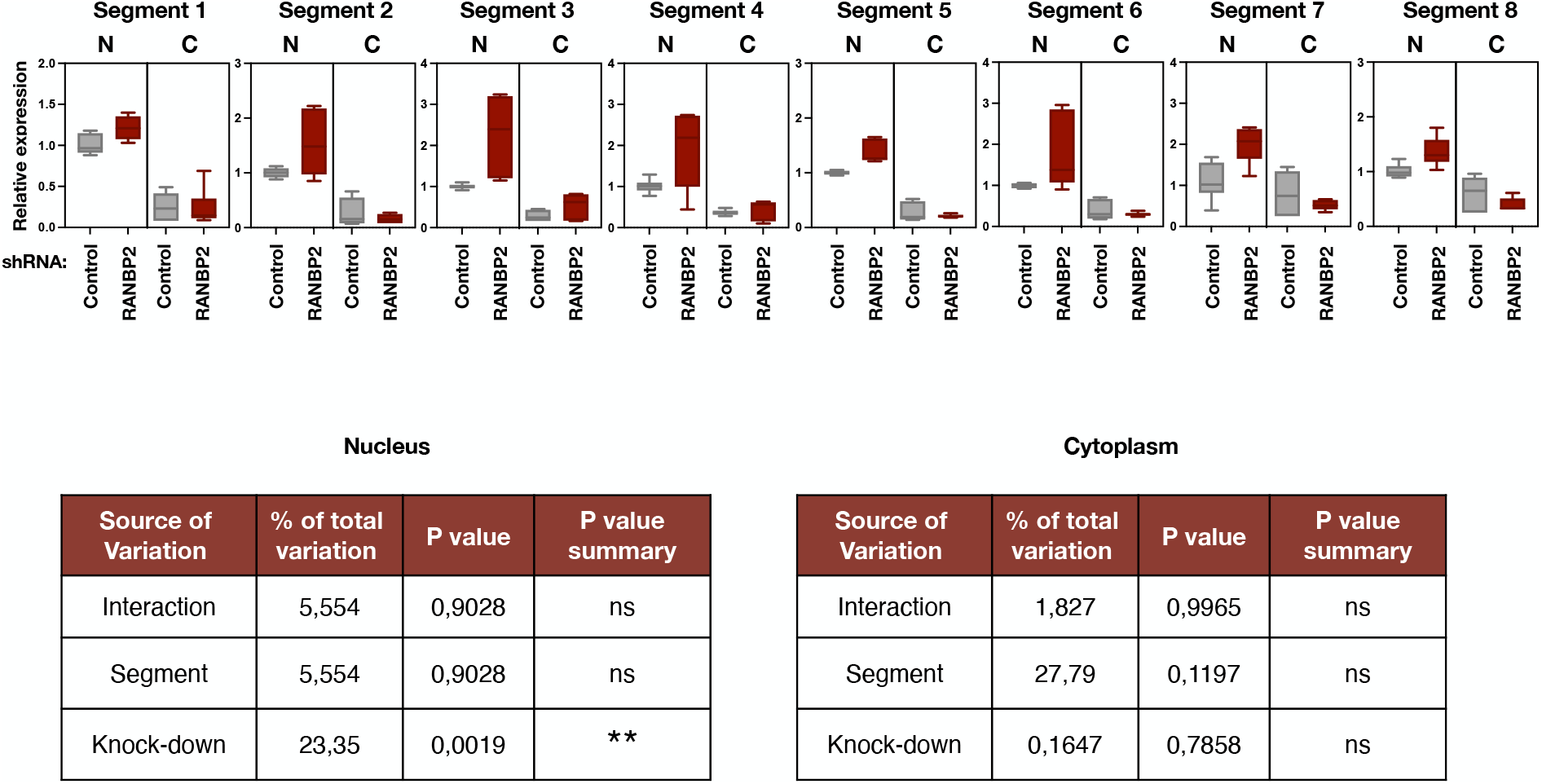
cRNA increase of all segments in the nucleus of RANBP2-KD cells. Transduced A549 cells were infected with IAV at MOI 0.5. At 6 hpi, nuclear and cytoplasmic fractions were separated and total intracellular RNAs were extracted. Strand-specific RT was performed with tagged primers to amplify complementary viral RNA (cRNA) specifically. Each of the 8 IAV segments was then amplified by qPCR using specific primers, as described in Figure 2G. Results include all technical replicates from n = 3 independent experiments, and are presented as mean +/− range, normalized to the nuclear fraction of the control condition. Two-way ANOVA (**P < 0.01, ns: non-significant) indicates that variations in the nuclear abundance of the cRNA segments varies significantly upon RANBP2 knockdown.

**Figure S10:**
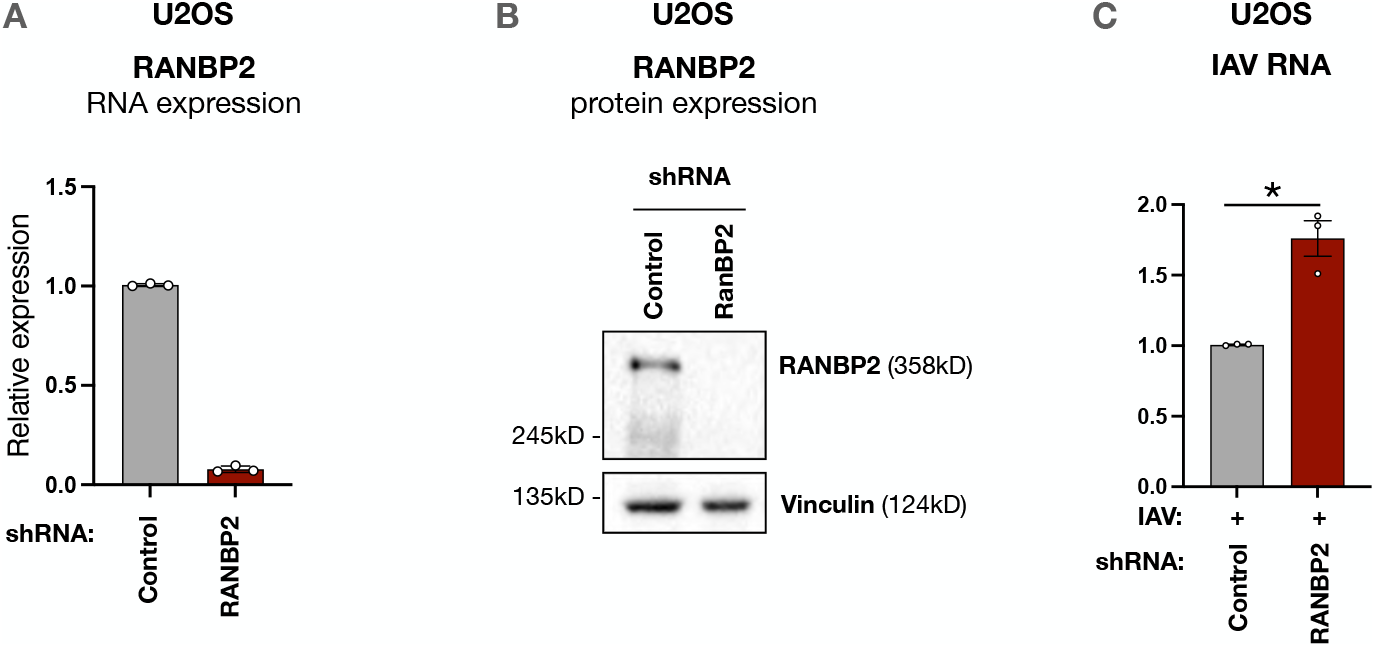
RANBP2 knockdown in U2OS cells leads to increased IAV RNA. Transduced A549 cells were infected with IAV at MOI 0.5. RANBP2 knockdown was quantified by **(A)** RTqPCR and **(B)** western blot, at 72 h post-transduction. **(C)** Total intracellular RNAs were extracted and viral RNAs encoding the Matrix 1 (M1) protein were amplified by RTqPCR. qPCR results (panels A, C) are normalized on RPL13a and to the control condition (mean +/− SEM). Each dot represents a biological replicate from n = 3 independent experiments. Two-tailed paired Student’s t test, *P < 0.05.

**Figure S11:**
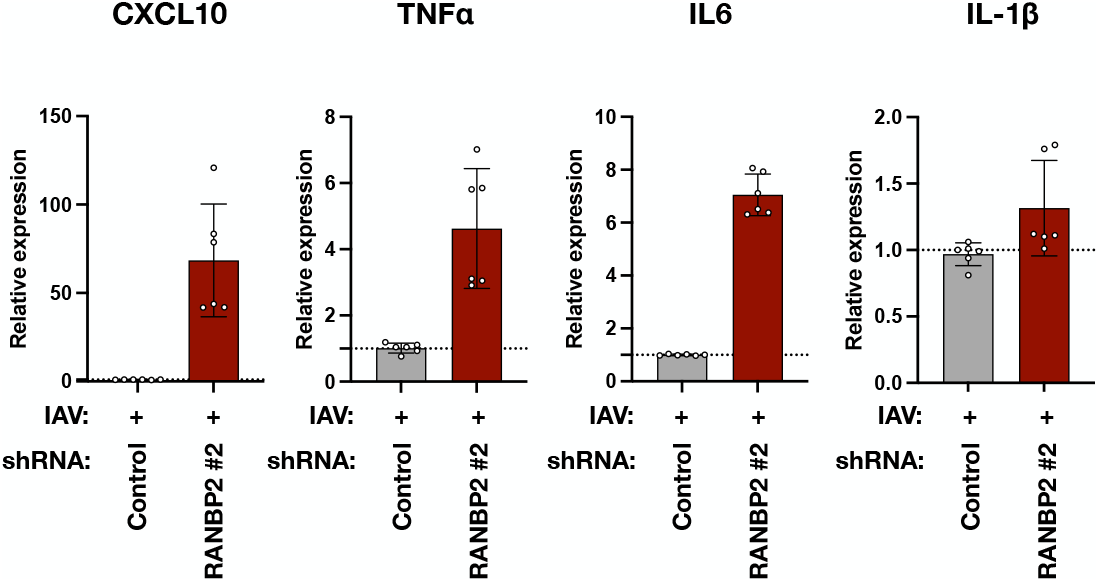
Increase in cytokine transcripts in RANBP2-KD cells, using a second shRNA. A549 cells were transduced with LV shRNA-control or shRNA-RANBP2#2 (see Figure S4A) for 48 h and infected with IAV (A/WSN/1933) at MOI 0.5. At 8 hpi, cytokine transcripts were assessed by RT-qPCR. Results were normalized on RPL13a and to the control condition (mean +/− SD).

**Figure S12:**
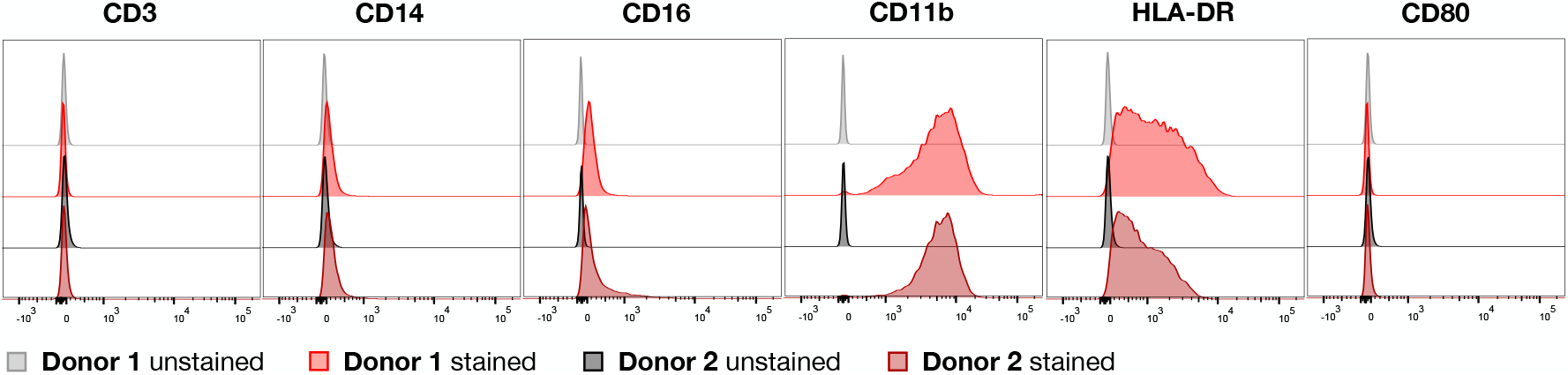
Phenotypic analysis of primary human MDM. PBMCs from two healthy donors were used to isolate monocytes by plastic adherence and differentiate them into macrophages by addition of GM-CSF. After 7 days of differentiation, MDMs were stained for phenotypic markers. The expression of CD3, CD14, CD16, CD11b, HLA-DR and CD80 was assessed by flow cytometry.

**Figure S13:**
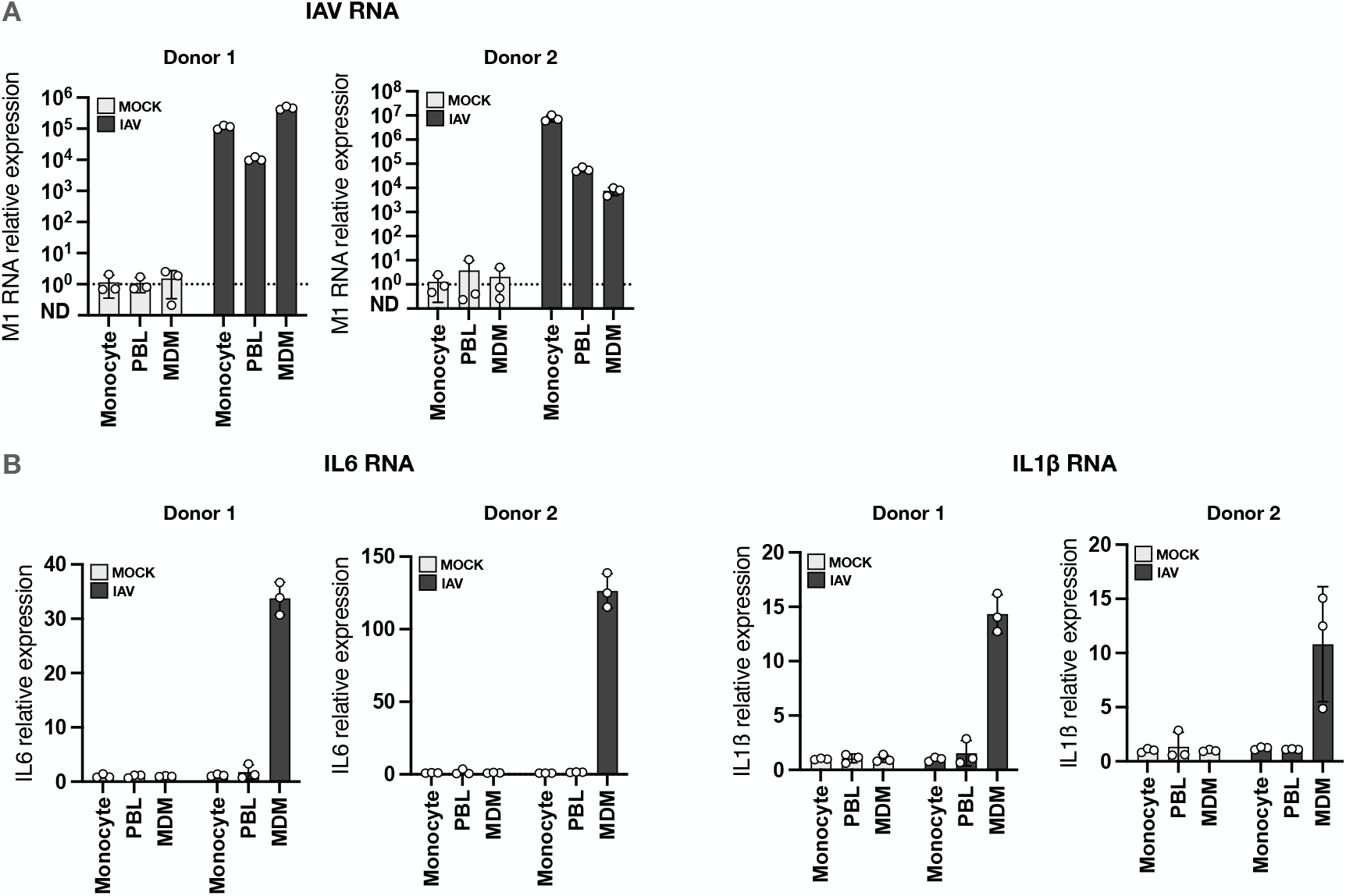
MDMs sense IAV infection by producing inflammatory cytokines (IL-6 and IL-1β) PBMCs from two healthy donors were used to isolate monocytes, peripheral blood lymphocytes (PBL), and MDMs. Cells were challenged with IAV (A/WSN/1933) at MOI 0.5 and total intracellular RNAs were extracted at 16 hpi. **(A)** M1-encoding viral RNAs and (**B)** IL-6 and IL-1β mRNAs were amplified by RTqPCR. Results are normalized on RPL13a and to the uninfected condition for each cell type represented, and are shown as mean +/− SD.

**Figure S14:**
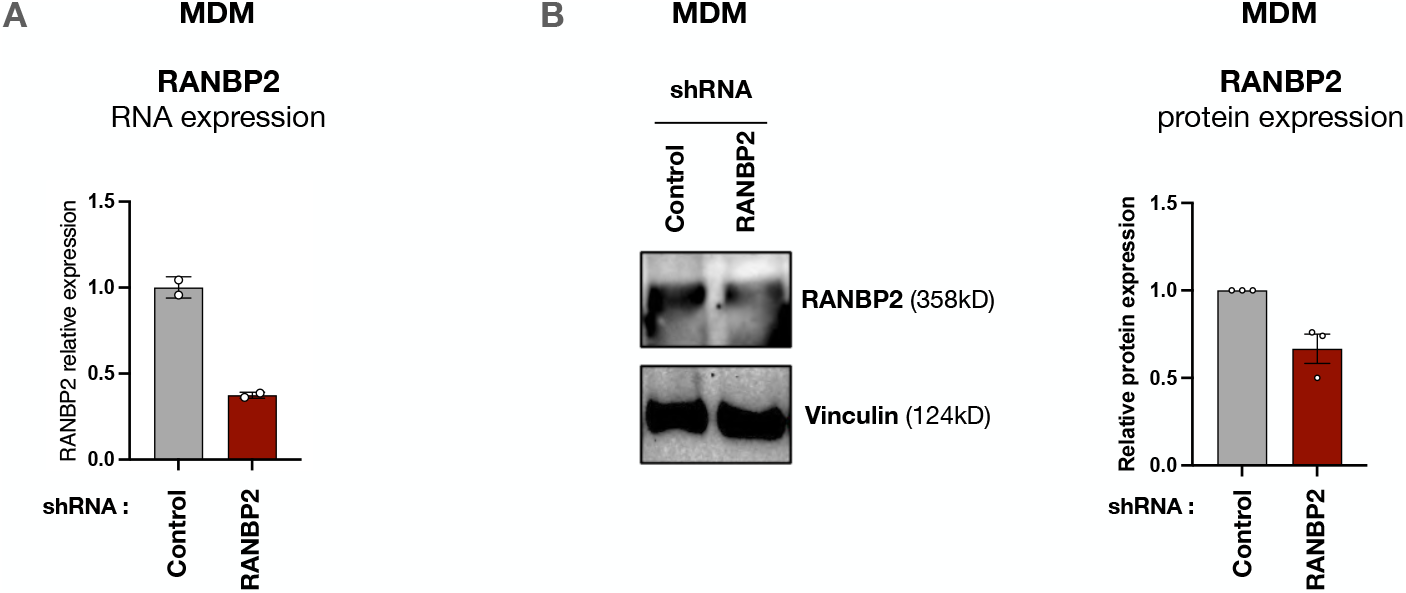
Assessment of RANBP2 knockdown in MDMs. MDM cells were transduced with LV shRNA-control or shRNA-RANBP2 at MOI 20 for 72 h. Knockdown efficacy was assessed by **(A)** RTqPCR and **(B)** western blot. Band intensities from n = 3 independent experiments were quantified using ImageLab. qPCR results (panel A) are normalized on RPL13a and to the control condition, while western blot quantifications (panel B) are normalized on the vinculin signal and to the control condition.

**Figure S15:**
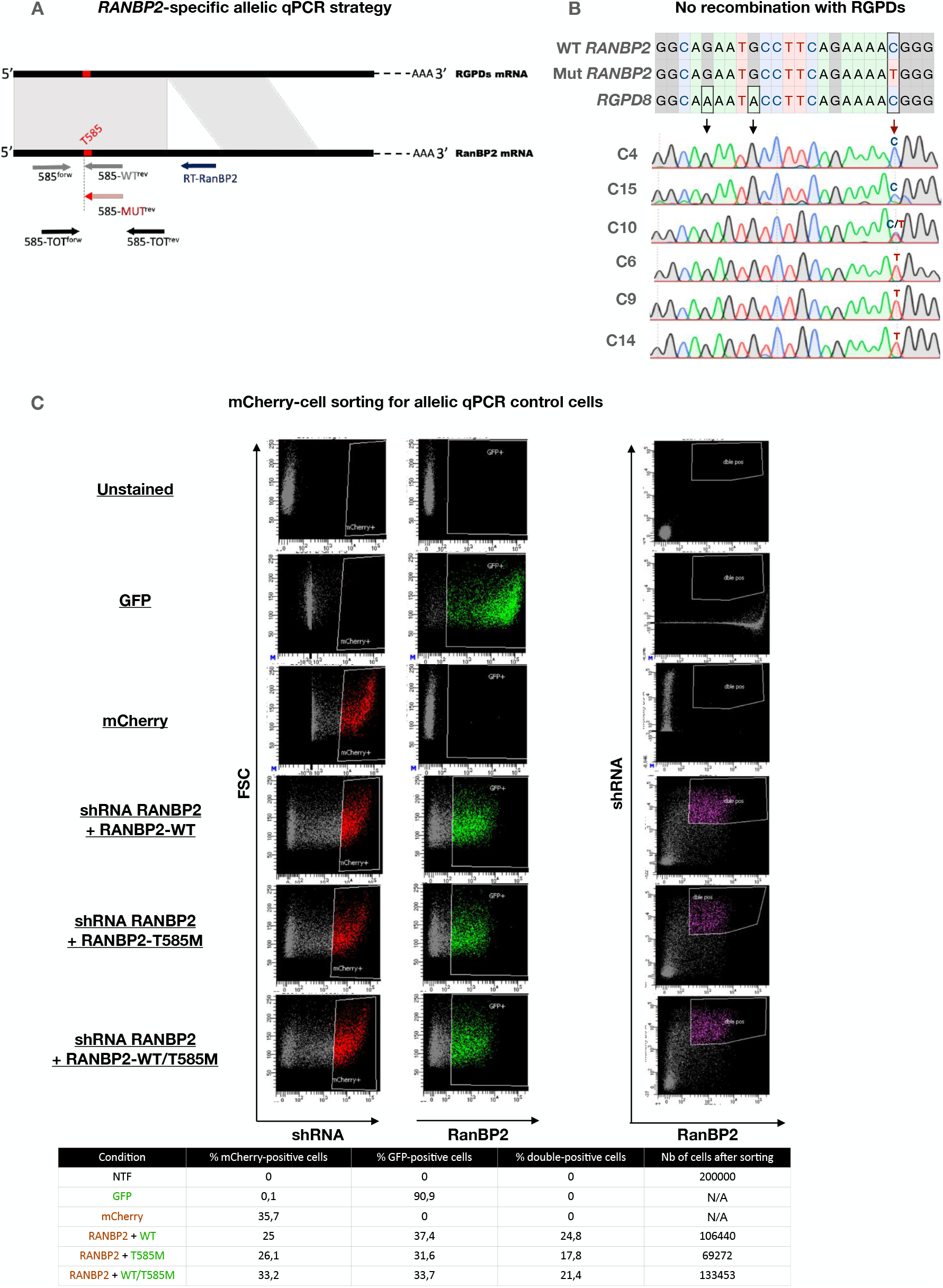
CRISPR-Cas9 knock-in strategy to generate ANE1-variants in U2OS cells. **(A)** RANBP2-specific allelic qPCR strategy. RANBP2-specific RT was performed using a primer unique to the RANBP2 sequence and absent in the highly homologous RGPD sequence (conserved regions are indicated in gray). Mutant and WT RANBP2 were distinguished using a reverse qPCR primer carrying either an A or a G at its 3’ end. A separate set of primers was also designed to amplify RANBP2 (near the same region) without distinguishing both forms. **(B)** Representation of the RANBP2 sequence upstream of the repair site, and one of the eight RGPDs. Sanger sequencing of the CRISPR clones shows no recombination with the RGPD sequence as all carry a G at the indicated positions (black arrows). **(C)** To produce controls for the screening of CRISPR clones by allelic qPCR, HEK-293T cells were transfected with mCherry-shRNA-RANBP2 and either GFP-RANBP2-WT, GFP-RANBP2-T585M, or both. mCherry-expressing cells were sorted by FACS and total intracellular RNAs were extracted for the allelic RTqPCR. Results of a representative FACS sorting are presented.

**Figure S16:**
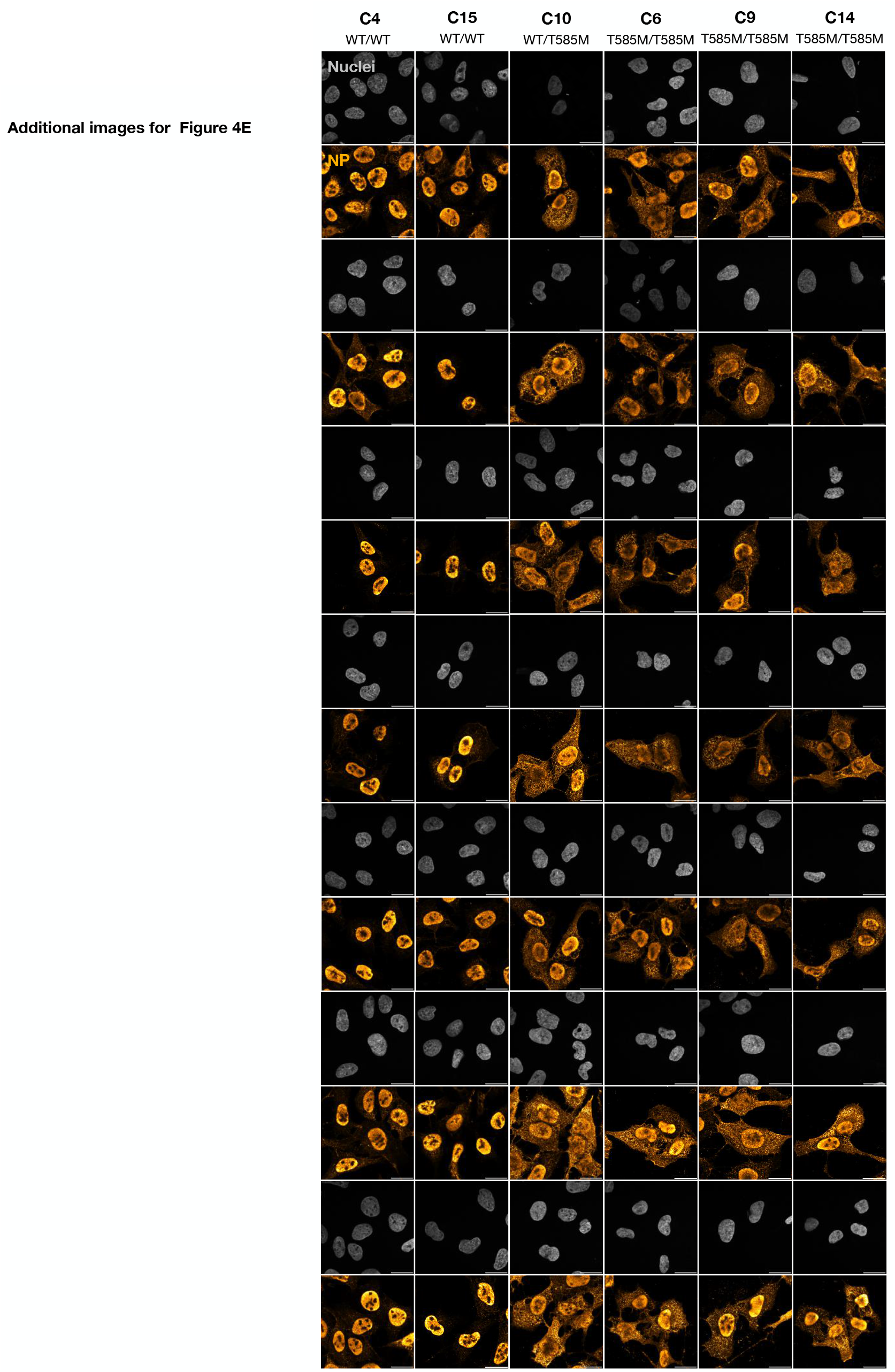
Increase in cytoplasmic NP in CRISPR RANBP2-T585M clones at 8 hpi. Full panel of images from n = 3 independent experiments for Figure 4E. CRISPR clones were infected with IAV (A/WSN/1933) at MOI 0.5. At 8 hpi, cells were fixed and stained for NP. Scale bar: 20µm. Image processing was performed using Fiji. NP (AlexaFluor-488) signal was pseudocoloured to “Orange Hot” for aesthetic purposes.

**Figure S17:**
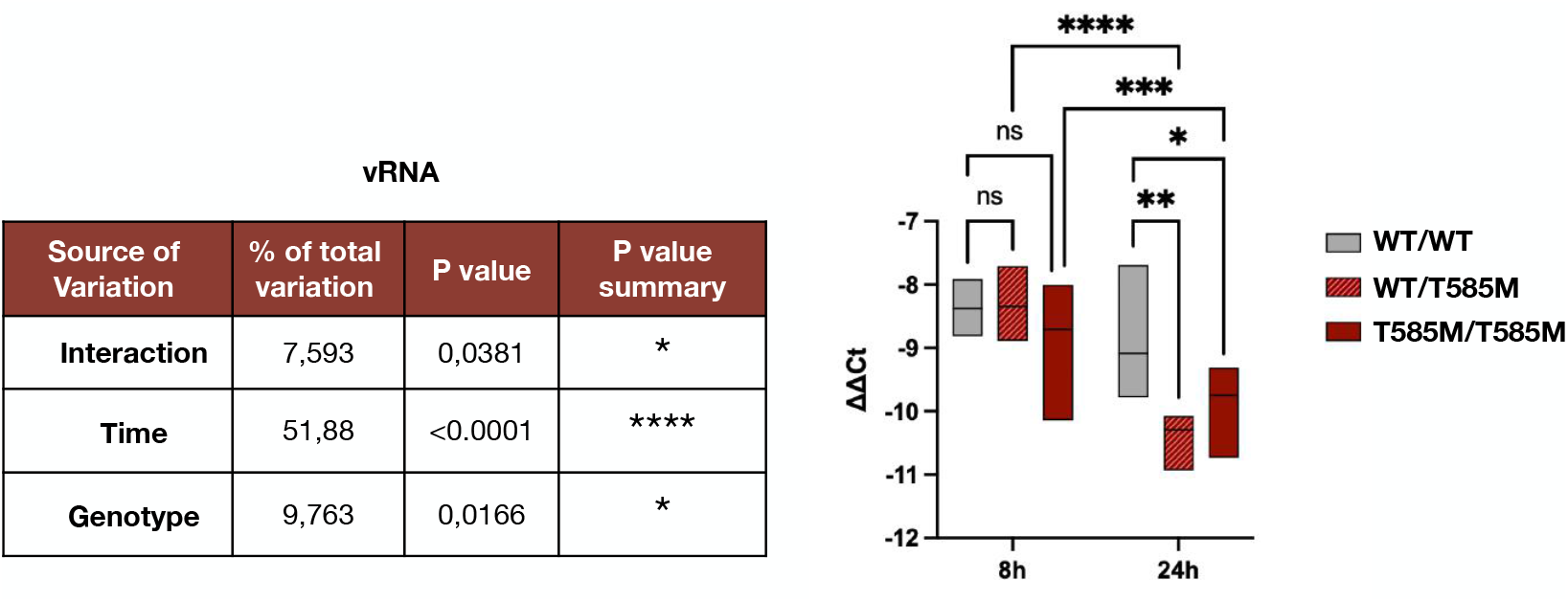
Increase in vRNA in CRISPR RANBP2-T585M clones. Statistical analysis of data presented in Figure 4G. qPCR vRNA signals at 8 and 24 hpi (MOI 0.5) were normalized to RPL13a (ΔCt), and to the corresponding early time point (1h) for each clone (ΔΔCt). Two-way ANOVA was performed on the mean ΔΔCt values from n = 3 independent biological replicates, with data pooled according to genotype groups (WT/WT, WT/T585M, and T585M/T585M). Statistical analysis revealed significant effects of both time and genotype, as well as a genotype × time interaction (p = 0.0381), indicating that the extent of vRNA accumulation over time differed among genotypes. At 24 hpi, both heterozygous and homozygous genotypes showed significantly higher vRNA levels compared to WT clones.

**Figure S18:**
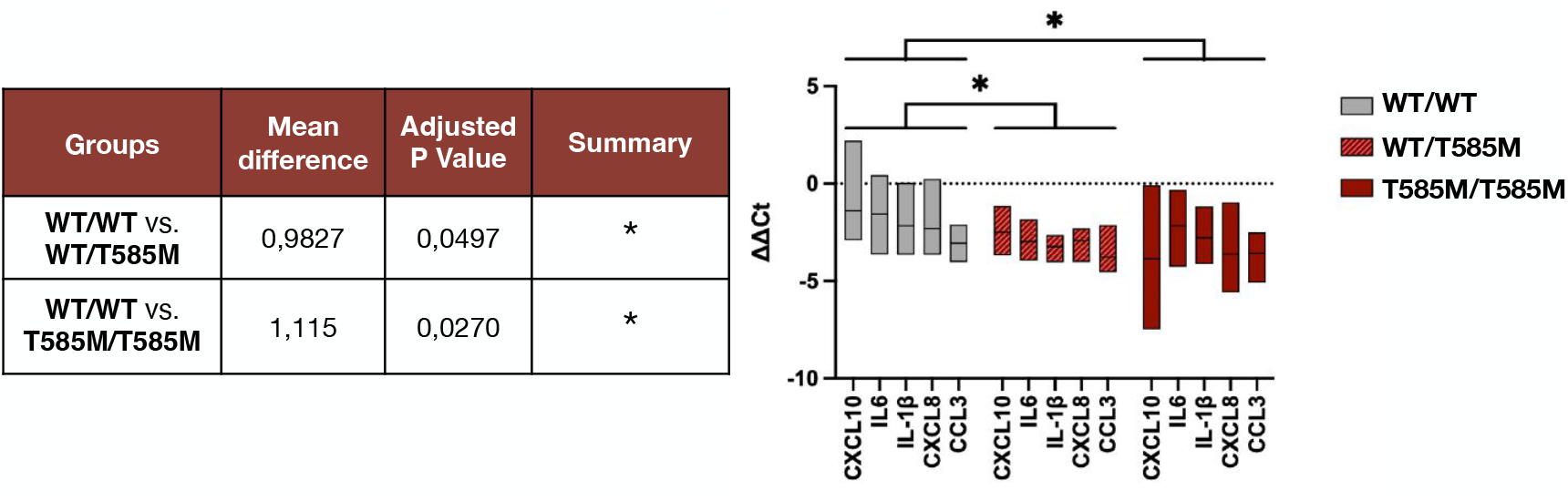
Increase in cytokine transcripts in CRISPR RANBP2-T585M clones. Statistical analysis of data presented in Figure 4H. qPCR cytokine transcript levels after overnight infection with IAV (MOI 0.5) were normalized to RPL13a (ΔCt) and to the corresponding uninfected condition for each clone (ΔΔCt). Ordinary one-way ANOVA was performed on the mean ΔΔCt values from n = 3 independent biological replicates, with data pooled according to genotype groups (WT/WT, WT/T585M, and T585M/T585M). Statistical analysis revealed significant overall differences in cytokine expression between WT clones and both heterozygous and homozygous ANE1 genotypes.

**Figure S19:**
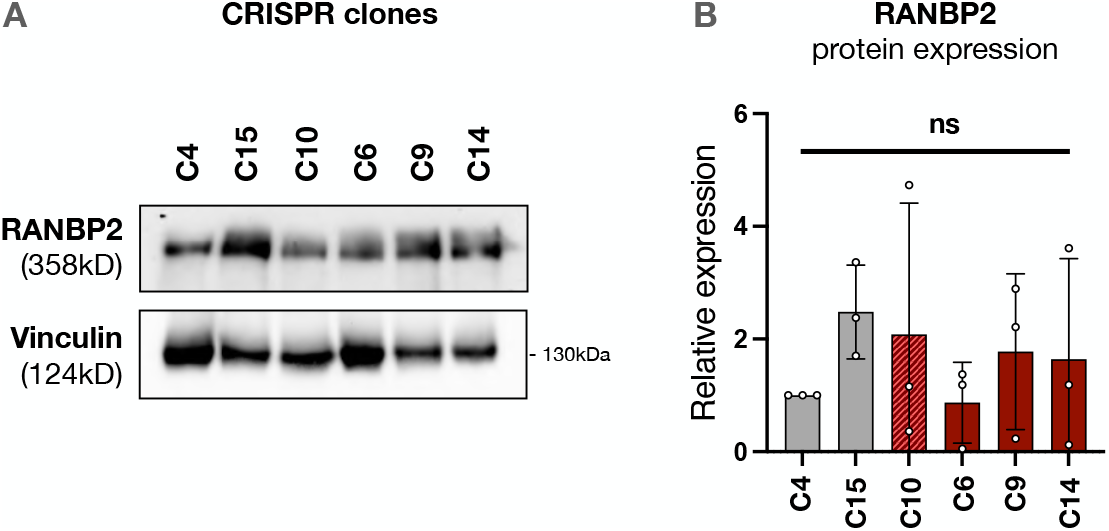
RANBP2 protein levels in CRISPR clones. **(A)** CRISPR clones were lysed and protein levels of RANBP2 were determined by western blot. **(B)** Band intensities from n = 3 independent experiments were quantified using ImageLab. Results are represented normalized to the vinculin signal and the C4 WT clone (mean +/− SD). One-way ANOVA. ns: non-significant.

## SUPPLEMENTARY TABLE LEGENDS

**Table S1: List of plasmids**.

**Table S2: List of reverse transcription and qPCR primers**.

**Table S3: List of antibodies used for immunofluorescence, flow cytometry and Western blotting**.

## Notes

### Competing Interest Statement

The authors have declared no competing interest.

### Summary of Updates

Following the reviewers' comments, we replicated the key experiments using alternative assays or shRNA, and we performed new experiments. In particular, to better characterize the RANBP2-T585M CRISPR clones, Figure 4 now includes evidence that the RANBP2-T585M point mutant increases IAV vRNA and innate immune signaling similarly to RANBP2 knockdown. Moreover, all datasets were reanalysed to ensure that statistical tests are performed correctly. In each case, mean values were first calculated from technical replicates, and statistical analyses were subsequently conducted using these mean values across biological replicates only, as requested.

