## Supplementary material for "The genetic driver of Acute Necrotizing Encephalopathy, *RANBP2*, regulates the inflammatory response to Influenza A virus infection": All Tables

**Table S1** **Plasmids**

| Target | Reference |
|---|---|
| pVSVg | pHCMV-G (Yee et al., PNAS 1994) |
| p8.74 | pCMVΔR8.74 (Dull et al., J Virol 1998) |
| mCherry-shRNA-Control | Addgene #1864 (Gouy et al., Front Neurol 2023) |
| mCherry-shRNA-RanBP2 | Gouy et al., Front Neurol 2023 |
| GFP-RanBP2-WT | Gift from J Joseph (Pune University) |
| GFP-RanBP2-T585M | This paper, availbale upon request |
| Centauri-Nluc-NLS | Fernandez et al., Proc Natl Acad Sci USA 2020 |

**Table S2** **Reverse transcription primers**

| IAV strand-specific RT | |
|---|---|
| Target | Sequence (5' -> 3') |
| vRNA | GGCCGTCATGGTGGCGAATAGCGAAAGCAGG |
| cRNA | CCAGATCGTTCGAGTCGTAGTAGAAACAAGG |

The colored sequence corresponds to the tag

| RanBP2-specific RT | |
|---|---|
| Target | Sequence (5' -> 3') |
| RanBP2 | ATGCTGTTGGGGTGGAAGCC |

**qPCR primers**

| Target | Forward primer (5'-3') | Reverse primer (5'-3') |
|---|---|---|
| CT-vRNA_Segment1 | CGAGACTGCGACATTAGATTCC | GGCCGTCATGGTGGCGAAT |
| CT-vRNA_Segment4 | GAGTTGTTCGCATGGTAGCC | GGCCGTCATGGTGGCGAAT |
| CT-cRNA_Segment1 | AGCGACCAAAAGAATTCGGATGG | CCAGATCGTTCGAGTCGT |
| CT-cRNA_Segment4 | GGGTCTTTGCAGTGCAGAATATGC | CCAGATCGTTCGAGTCGT |
| IAV M1 | CAGCACTGGAGCTAGGATGA | AAGGCTATGGAGCAAATGGC |
| IAV NP | CCTCGGGCCAAATCAGCATA | ATTCCCAGTGAATGCTGCCA |
| Segment 1 | CGGGATATTGCCCGACATGA | TCCGCGCTGGAATACTCATC |
| Segment 2 | TACCGGTGCCATAGAGGTGA | TGGGTTTGCTCCCACAGTTT |
| Segment 3 | CAAGCTTGCCGACCAAAGTC | TTGCCCTCAATGTAGCCGTT |
| Segment 4 | GGCATCATCACCTCAAACGC | CCCTGGGGTGTTTGACACTT |
| Segment 5 | TCAAGTGAGAGAGAGCCGGA | TGAGTGCAGACCGTGCTAAA |
| Segment 6 | TTAATGAGCTGCCCTGTCGG | GCTGACCAAGCAACCGATTC |
| Segment 7 | CCACTGAAGTGGCATTTGGC | GCTGGGAGTCAGCAATCTGT |
| Segment 8 | GCTCATGCCCAAGCAGAAAG | CCATGATCGCCTGGTCCATT |
| RanBP2 (Total) | AAAACATGGCCTTCAACCTG | TCAACAATTTCTGATGCCTGA |
| RanBP2 (WT) | TGGGATGCGGTTTGTACTCT | AGAATTAAGACCGCTGCCCG |
| RanBP2 (T585M) | TGGGATGCGGTTTGTACTCT | AGAATTAAGACCGCTGCCCA |
| RanBP2 (Exon24) | CAGGAAGCTCAAAAATCTCAGACA | ACCATCACTTCAGTCCCACC |
| IL-6 | TAACCACCCCTGACCCAACC | ATTTGCCGAAGAGCCCTCAG |
| IL-1ß | GGCATCCAGCTACGAATCTC | GAACCAGCATCTTCCTCAGC |
| TNFα | GGCGTGGAGCTGAGAGATAAC | GGTGTGGGTGAGGAGCACAT |
| IL-18 | GGAATTGTCTCCCAGTGCAT | ACTGGTTCAGCAGCCATCTT |
| CXCL10 | GTCCACGTGTTGAGATCATTGC | TTGCTCCCCTCTGGTTTTAAGG |
| RPL13a | AACAGCTCATGAGGCTACGG | TGGGTCTTGAGGACCTCTGT |
| Actine B | CTGGAACGGTGAAGGTGACA | AAGGGACTTCCTGTAACAATGCA |

**Table S3** **Antibodies**

| Immunofluorescence | | | |
|---|---|---|---|
| Target | Reference | Brand | Dilution |
| NP | MCA400 | Biorad | 1/1000 |
| PB1 | GTX125923 | GeneTex | 1/500 |
| RANBP2 | 27606-1-AP | Proteintech | 1/100 |
| FG repeats NPC (clone mab414) | 902901 | Biolegend | 1/200 |
| Goat anti-Mouse AF488 | A11001 | Thermofisher Scientific | 1/1000 |
| Goat anti-Rabbit AF488 | A11034 | Thermofisher Scientific | 1/1000 |
| Goat anti-Mouse AF555 | A21424 | Thermofisher Scientific | 1/1000 |

| Flow cytometry and MDM phenotyping | | | |
|---|---|---|---|
| Target | Reference | Brand | Dilution |
| HA (Centauri) | 11988506001 | Roche, Sigma | 1/100 |
| CD3-BV421 (clone UCHT1 | 300433 | Biolegend | 1/100 |
| CD14-PerCP-Cy5.5 (clone HCD14) | 325621 | Biolegend | 1/100 |
| CD16-Alexa700 (clone 3G8) | 302026 | Biolegend | 1/100 |
| CD11b-APC-Cy7 (clone M1/70) | 101225 | Biolegend | 1/100 |
| HLA-DR-FITC (REA805) | 130-111-941 | Miltenyi | 1/100 |
| CD80-BV650 (clone 2D10) | 305227 | Biolegend | 1/100 |

| Western Blot | | | |
|---|---|---|---|
| Target | Reference | Brand | Dilution |
| NP | MCA400 | Biorad | 1/5000 |
| PB1 | GTX125923 | GeneTex | 1/5000 |
| PB2 | GTX125925 | GeneTex | 1/5000 |
| PA | GTX125932 | GeneTex | 1/5000 |
| Actin ß | A1978 | Roche, Sigma | 1/3000 |
| Lamin A/C | 10298-1-AP | Proteintech | 1/400 |
| Tubulin | ab6160 | Abcam | 1/5000 |
| Vinculin | 26520-1-AP | Proteintech | 1/5000 |
| RANBP2 | PA1-O82 | Invitrogen | 1/500 |
| Goat anti-Mouse HRP | NA931V | GE Healthcare | 1/5000 |
| Donkey anti-Rabbit HRP | NA934V | GE Healthcare | 1/5000 |
| Goat anti-Rat HRP | ab97057 | Abcam | 1/5000 |
